# Butyrate preserves entorhinal-hippocampal spatial coding and blood brain barrier integrity in mice with depleted gut microbiome

**DOI:** 10.1101/2025.07.24.666609

**Authors:** Joshua M. Glynn, Joshua J. Strohl, Ciara Bagnall-Moreau, Joseph Carrión, Patricio T. Huerta

## Abstract

Given the widespread and increasing consumption of oral antibiotics globally, understanding their impact on cognition through the gut-brain axis is crucial. We investigated whether broad-spectrum antibiotics disrupt spatial cognition by altering behavior, neural dynamics, brain metabolism, and blood-brain barrier integrity. Here we show that male mice receiving antibiotic-treated water display significant impairments in spatial memory tasks and abnormal encoding of space by entorhinal grid cells and hippocampal place cells. These cognitive deficits are accompanied by altered brain metabolism and blood-brain barrier permeability in the hippocampal formation. Remarkably, supplementation with butyrate, a key microbiome-derived metabolite, preserves spatial cognition, neural dynamics, and blood-brain barrier function despite antibiotic treatment. These findings reveal that gut microbiome depletion disrupts the hippocampal-entorhinal network underlying spatial cognition, while suggesting butyrate supplementation as a potential therapeutic approach to mitigate antibiotic-induced cognitive impairments.

## INTRODUCTION

Antibiotics have been tremendously beneficial in reducing mortality by protecting against infection, but their widespread use has raised concerns about adverse effects. In the United States alone, 236.4 million antibiotic prescriptions were issued in 2022 (data from the Centers for Disease Control and Prevention), with at least 30% deemed inappropriate for the conditions being treated (*1*, *2*). Global antibiotic consumption has increased by at least 40%, particularly in low- and middle-income countries across South Asia, Eastern Europe, North Africa, and the Middle East (*3*, *4*). This excessive use has contributed to rising antimicrobial resistance (AMR), which caused an estimated 1.27 million deaths in 2019 (*5*, *6*). Because of AMR, the utility of antibiotic combination therapies in combating resistant infections has been intensely studied in both the clinic and in vitro (*7*, *8*). However, there is a growing concern that the consumption of oral antibiotics can lead to substantial abnormalities in the gut microbiome in humans and animals (*9–12*). Additionally, it is clear that oral antibiotics disrupt the gut-brain axis, as rodents treated with antibiotics exhibit abnormal behaviors related to addiction, social interaction, memory, and anxiety (*13–18*). These tantalizing observations open the window to study how the gut microbiome affects the brain centers that underlie cognition. In particular, the potential interaction between the microbiome and the neural substrates for spatial cognition, such as the hippocampus and the entorhinal cortex (*19–21*), remains largely unexplored.

The hippocampus contains place cells, which are pyramidal neurons that exhibit localized firing at specific locations within a defined space. Place cells generate place fields that span the area of an environment, thereby creating a representation of space in the brain (*21–26*). The entorhinal cortex modulates place cell encoding through direct inputs, as lesions to this area cause place field expansion (*27*). Additionally, the entorhinal cortex contains grid cells that, like place cells, fire in distinct locations but uniquely tessellate space in equilateral triangular patterns (*28*, *29*). Disruption of this hippocampal-entorhinal network in various disease states alters both place and grid cell dynamics and impairs spatial memory (*30–32*). This creates a tangible framework to investigate how antibiotic-induced gut microbiome depletion affects spatial cognition, in the context of behavior and neural dynamics.

Animals receiving oral antibiotics develop a markedly altered metabolome that likely contributes to cognitive impairment through multiple facets of the gut-brain axis. Antibiotic treatment particularly affects short chain fatty acids (SCFAs), which are produced in abundant quantities by over 200 species of the gut microbiome primarily through fermentation of dietary fibers (*33*, *34*). SCFAs help maintain the integrity of both the blood-brain barrier (BBB) and the intestinal epithelial barrier by inducing the expression of tight junction proteins (*35–37*). Gut microbiome depletion reduces SCFA levels in the cecum and feces, downregulates tight junction proteins, and compromises barrier integrity, resulting in both ‘leaky gut’ and ‘leaky brain’ conditions (36, *38*–*42*). Furthermore, ex vivo and in vivo studies using anesthetized animals suggest that depletion of the gut microbiome perturbs synaptic plasticity within the hippocampus (*43–45*). However, there is still a limited understanding of how spatial cognition and the corresponding neural substrates are affected in actively behaving animals with a depleted gut microbiome. This study demonstrates that antibiotic treatment impairs spatial cognition and disrupts spatial encoding within both the hippocampus and entorhinal cortex of freely behaving mice. We complement these findings with brain positron emission tomography (PET) showing altered brain metabolism and BBB permeability. Importantly, we describe that supplementation with butyrate, a SCFA secreted by the gut microbiome, is sufficient to preserve spatial memory, the integrity of CA1 place cells, and BBB function despite gut microbiome elimination.

## RESULTS

### Antibiotic treatment leads to impaired spatial cognition

We investigated whether antibiotics affect spatial cognition by treating C57BL/6 mice with a broad-spectrum antibiotic cocktail, consisting of ampicillin, neomycin, gentamicin, metronidazole, and vancomycin in their drinking water (termed ABX mice henceforth) (**Fig. 1A**). Control mice (termed CON henceforth) drank regular water. The ABX group started receiving antibiotics two weeks prior to behavioral testing and was maintained on this treatment throughout experiments (**Fig. 1A**). Notably, ABX mice exhibited impaired spatial cognition in the clockmaze task, as evidenced by significantly longer escape latencies and greater distances traveled to locate the true exit, despite making a similar number of errors compared to CON mice (**Fig. 1B**). For each trial, we classified the strategy used by an animal as futile (did not find the true exit in <60s), chain (found the true exit in <60s, made >4 errors) or spatial (found the true exit in <60s, made ≤4 errors). ABX mice consistently failed to use the spatial strategy across trials (**Fig. 1C**), which we further quantified with a strategy score (see Methods for details) that was significantly lower in ABX mice compared to the CON group (**Fig. 1C**). After completing 12 trials, mice were subjected to a probe test (60s with all exits blocked). Unlike CON mice, which spent significantly more time in the target zone (where the true exit was formerly located), ABX mice showed no preference between target and non-target zones (fig. S1A). In the object-place memory (OPM) task, both CON and ABX mice spent similar amounts of time investigating the objects in the sample and choice phases of the task, however ABX mice had significantly lower OPM ratios compared to CON mice (**Fig. 1D**). We performed further behavioral tasks including the novel object recognition task (**Fig. 1E**), rotarod (fig. 1SB), elevated plus maze (fig. 1SC), light-dark chamber (fig. 1SD), and zero maze tasks (fig. 1SE). In contrast to our spatial cognition findings, we did not find any significant differences between CON and ABX mice in any of these tasks.

**Fig. 1.**
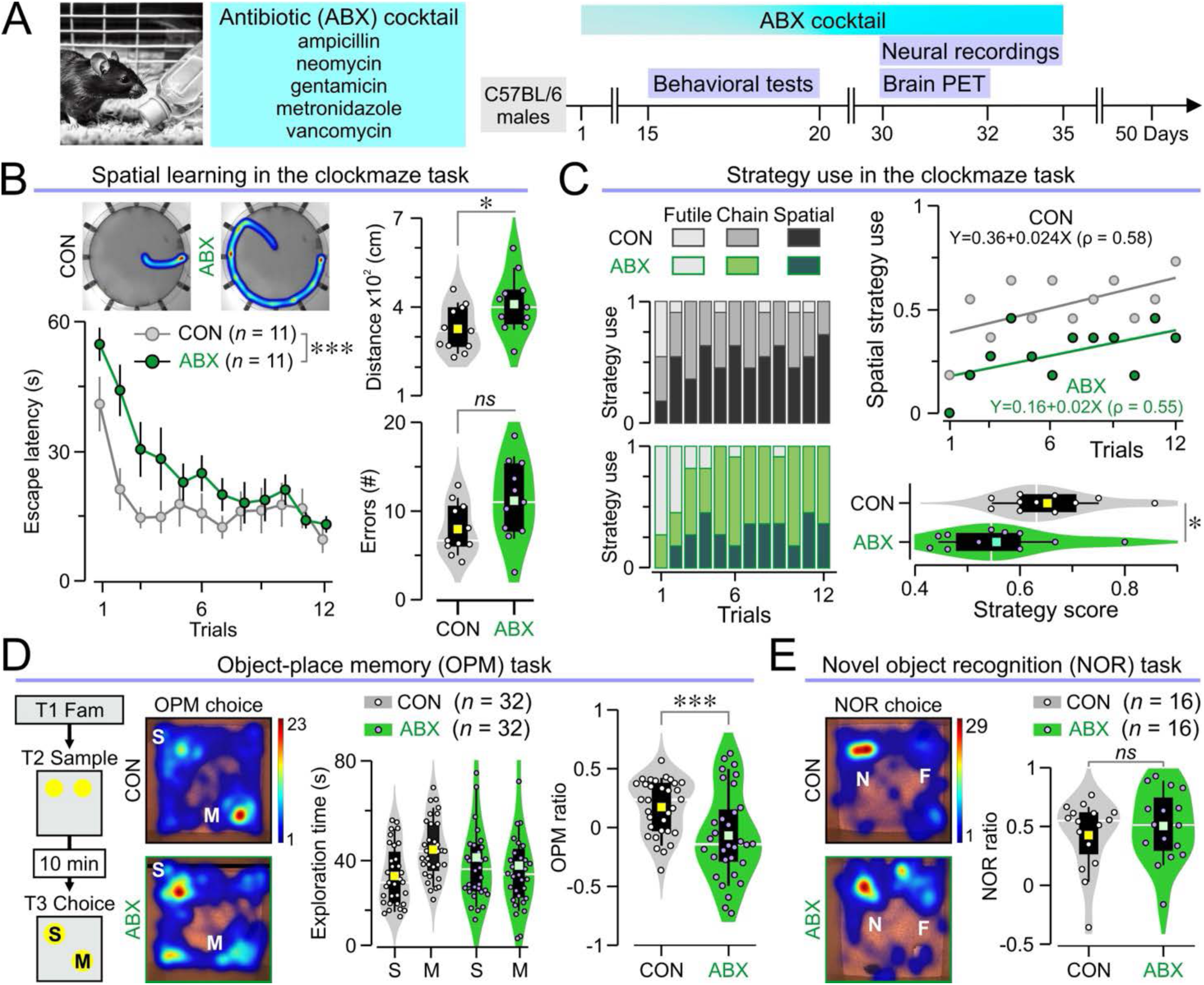
ABX mice display impaired spatial cognition. **(A)** Timeline of experimental procedures in mice given antibiotic (ABX) cocktail in the drinking water. Control (CON) mice drink regular water. (**B)** *Left,* heatmaps of representative paths traveled by mice in the clockmaze (*top*) and time series of escape latency (mean ± SEM) showing that ABX mice are significantly slower across trials in the clockmaze task (\**P* = 0.012, RMANOVA of log10 transformed times followed by Bonferroni test; *n* = 11 CON, 11 ABX). *Right,* violin plots showing average distances traveled (*top*, \**P* = 0.038, *t* test; *n* = 11 CON, 11 ABX) and number of errors made in all trials (*bottom*, *P* = 0.059, *t* test; *n* = 11 CON, 11 ABX). (**C)** ABX mice use less spatial strategy in the clockmaze, as revealed by the stacked column graphs showing the frequency of the different strategies (*left*). This is further demonstrated by the scatterplot of spatial strategy use (fitted with linear regression curves) for both groups (*top right*) and the violin plots showing strategy scores (*bottom right,* \**P* = 0.038, *t* test; *n* = 11 CON, 11 ABX) in which higher values reflect higher spatial use. (**D)** *Left,* schematic of the OPM task and representative occupancy heatmaps. *Middle,* violin plots showing the investigation times for the stable and moved objects (CON v. ABX, *P* = 0.866; stable v. moved, *P* = 0.213, two-way ANOVA; *n* = 32 CON, 32 ABX). *Right,* violin plots of the OPM ratios (\*\*\**P* = 0.003, *t* test; *n* = 32 CON, 32 ABX). (**E)** *Left,* representative occupancy heatmaps for the NOR task. *Right,* violin plots of the NOR ratios (*P* = 0.534, MWU test; *n* = 16 CON, 16 ABX).

### Antibiotic treatment disrupts brain glucose metabolism

To assess the regional effects of antibiotic treatment on brain metabolism, we injected mice intraperitoneally (IP) with [¹⁸F]fluorodeoxyglucose (FDG). Mice were scanned 40 min later, as this time window captures the maximal glucose uptake in the brain. We performed a brain-wide analysis to compare mean FDG signals between groups at the voxel level, using a stringent cutoff of *P* < 0.001 for significance. We identified bilateral clusters exhibiting significantly lower FDG uptake in the ABX group, which primarily encompassed the entorhinal cortex, but also included parts of the ventral CA1 field of the hippocampus and the subiculum. Given their symmetry, we quantified these bilateral clusters as a single unit, which showed significantly lower FDG standard uptake values (SUVs) in ABX mice compared to CON mice (**Fig. 2A**). At *P* < 0.05, these clusters expanded into neighboring regions, revealing a new set in the dorsal CA1 field of the hippocampus (**Fig. 2A**). Using the strict *P* < 0.001 cutoff, we also identified clusters of increased FDG uptake in the anterior hypothalamus and the striatum. We quantified the uptake for the cluster overlaying the anterior hypothalamus and found it to be significantly higher in ABX mice (**Fig. 2B**). Decreasing the threshold for significance revealed an expansion of the clusters into the striatum, bilaterally. Together, these results indicate that antibiotic treatment alters brain glucose metabolism, resulting in decreased uptake within the hippocampal formation (particularly the entorhinal cortex) and increased uptake in the anterior hypothalamus and striatum.

**Fig. 2.**
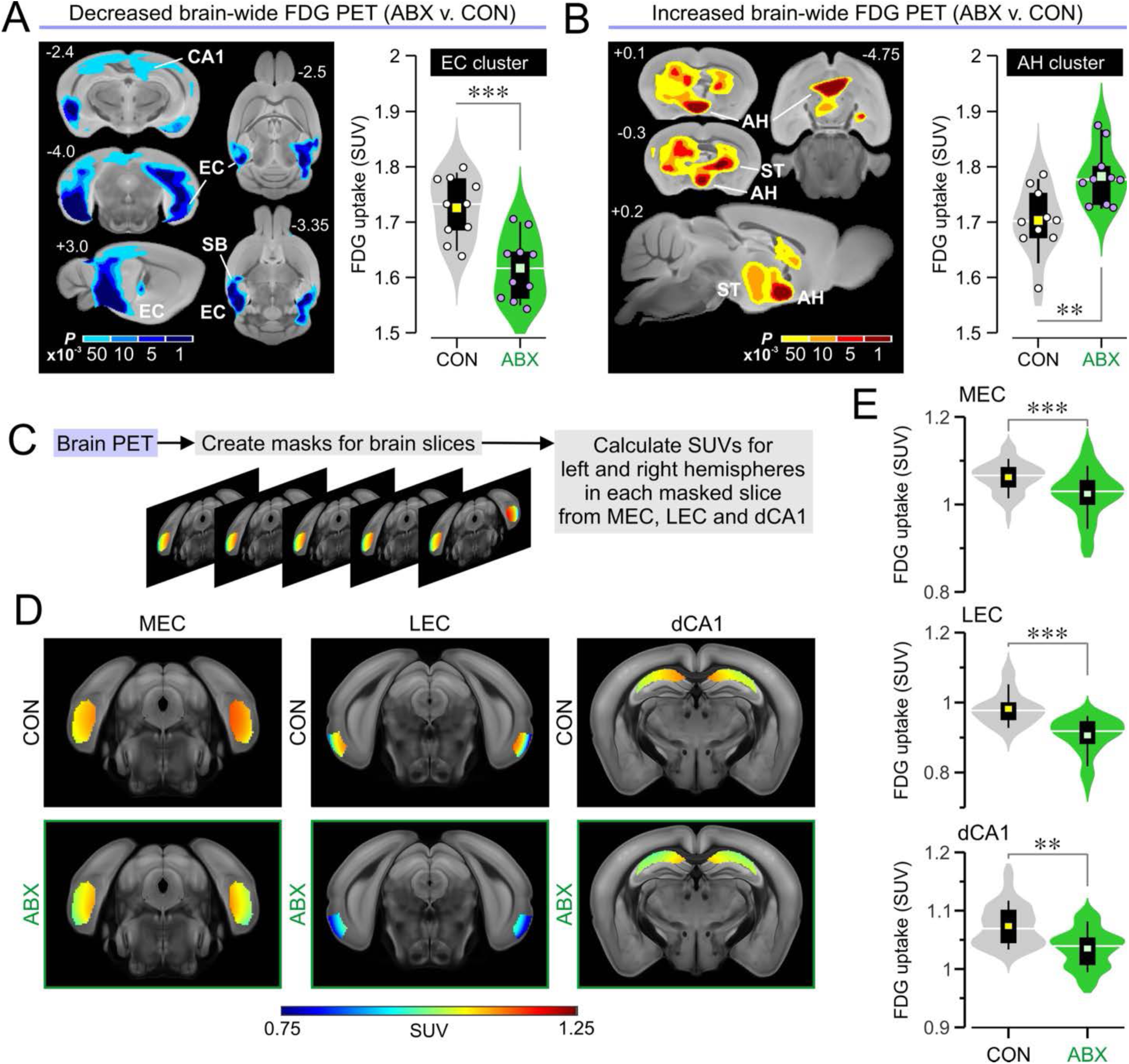
Positron emission tomography reveals disrupted brain glucose metabolism in ABX mice. **(A)** *Left,* data for voxel- wide search over the whole brain showing clusters in which FDG standard uptake units (SUVs) are significantly decreased in ABX mice when compared to CON mice, using increasing thresholds of significance (scale bar for *P* values <0.001, <0.005, <0.01, <0.05); abbreviations: CA1, CA1 field of the hippocampus; EC, entorhinal cortex; SB, subiculum. *Right,* violin plots showing FDG SUVs for the EC cluster (\*\*\**P* < 0.001, *t* test; *n* = 10 CON, 10 ABX). **(B)** *Left,* brain-wide data for clusters with significantly increased FDG SUV in ABX mice; abbreviations: AH, anterior hypothalamus; ST, striatum. *Right,* violin plots showing FDG SUVs for the AH cluster (\*\**P* = 0.005, *t* test; *n* = 10 CON, 10 ABX). (**C)** Schematic depicting the approach to brain atlas-based analysis in which masked slices are selected in the medial entorhinal cortex (MEC), lateral entorhinal cortex (LEC) and dorsal CA1 (dCA1). (**D)** Heatmaps showing the SUVs for representative slices of MEC, LEC, and dCA1 (each slice shows the averaged SUV across all mice in the group) overlaid onto an MRI template. (**E)** Violin plots showing that ABX mice display significantly decreased FDG uptake in the MEC (\*\*\**P* = 0.006, MMANOVA; *n* = 300 slices from 10 CON, 300 slices from 10 ABX), LEC (\*\*\**P* < 0.001, MMANOVA; *n* = 200 slices from 10 CON, 200 slices from 10 ABX), and dCA1 (\*\**P* = 0.009, MMANOVA; *n* = 300 slices from 10 CON, 300 slices from 10 ABX).

Our group has developed a novel atlas-based approach to analyze PET signals in precise anatomically defined brain regions (*46*). To specifically examine the neural substrates of spatial cognition (**Fig. 2C**), we measured FDG uptake in the medial entorhinal cortex (MEC), the lateral entorhinal cortex, and the CA1 region of the dorsal hippocampus. This atlas-based analysis revealed significantly decreased FDG SUVs in all three areas (**Fig. 2, D and E**).

### Impaired grid cell networks in the MEC of ABX mice

We sought to determine the neural substrate for impaired spatial cognition in ABX mice. Guided by the strong decrease in glucose metabolism in the MEC, we implanted multielectrode arrays into the MEC of CON and ABX mice for behavioral neurophysiology recordings (**Fig. 3A**) which yielded single unit activity arising from different neuronal types. To classify these units, we established a novel unsupervised machine learning pipeline. After feature extraction and normalization, we performed dimensionality reduction with uniform manifold approximation and projection (UMAP), which was followed by density-based spatial clustering of applications with noise (DBSCAN) (fig. S2, A and B; see Methods for details). For each cluster, we obtained z- scored spatial information (SI), grid score, border score, and mean firing rate (MFR). We classified cluster 1 as interneurons (high MFR, low SI), cluster 2 –which was the most populous– as grid cells (high grid scores), cluster 3 as undefined (few units, low border scores, low MFRs), cluster 4 as aperiodic spatial cells (high SI, low grid scores), and cluster 5 as border cells (high border scores) (**Fig. 3B** and fig. S2, C and D).

**Fig. 3.**
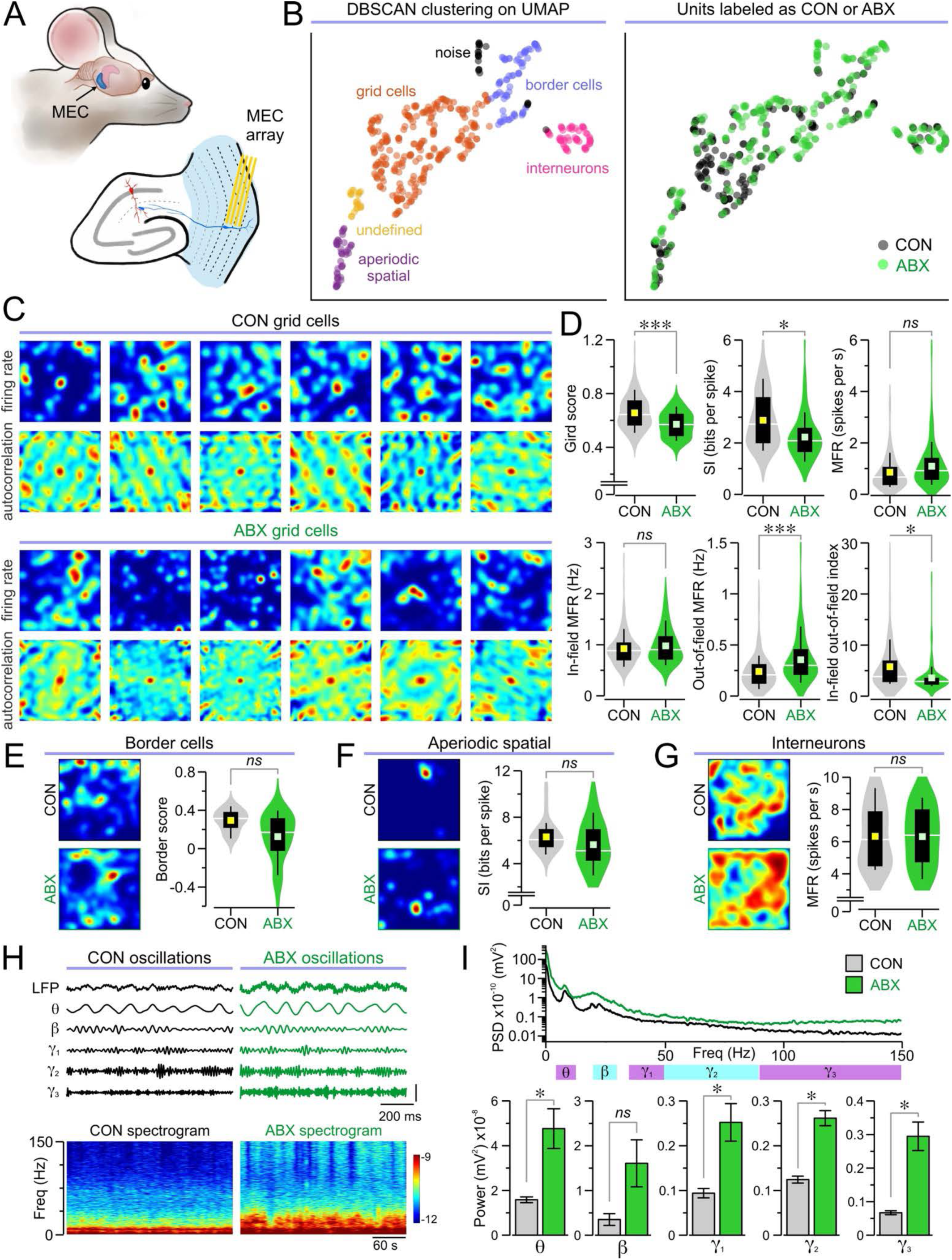
Neurophysiological recordings in the medial entorhinal cortex. **(A)** Illustration depicting the position of the tetrode array placement in the MEC. (**B)** *Left,* UMAP with DBSCAN clustering identifies interneurons, grid cells, undefined cells, aperiodic spatial cells, border cells, and noise. *Right,* UMAP showing the group (CON or ABX) for each unit. **(C–D)** Analysis of grid cells demonstrates large disruptions of grid cell properties in ABX mice compared to CON mice. (**C)** Representative grid cell rate maps and corresponding autocorrelations in both groups. (**D)** Violin plots of the grid score (\*\*\**P* < 0.001, MMANOVA), spatial information (SI, \**P* = 0.028, MMANOVA), mean firing rate (MFR, *P* = 0.057, MMANOVA), in- field MFR (*P* = 0.257, MMANOVA), out-of-field MFR (\*\**P* = 0.001, MMANOVA), and in-field-out-of-field index (\*\*\**P* < 0.001, MMANOVA) for each grid cell (*n* = 135 cells from 4 CON, 117 cells from 4 ABX). **(E–G)** Other cell types are similar between ABX and CON groups. (**E)** *Left,* representative border cell rate maps. *Right,* violin plots of border score (*P* = 0.147, MMANOVA; *n* = 18 cells from 3 CON, 55 cells from 4 ABX) for each border cell. (**F)** *Left,* sample aperiodic spatial cell rate maps. *Right,* violin plots of the SI (*P* = 0.204, MMANOVA; *n* = 24 cells from 4 CON, 20 cells from 3 ABX) for each aperiodic spatial cell. (**G)** *Left,* representative interneuron rate maps. *Right*, violin plots of the MFR (*P* = 0.941, MMANOVA; *n* = 19 cells from 4 CON, 37 cells from 4 ABX) for each interneuron. **(H–I)** Analysis of local field potentials (LFPs) during movement bouts reveal much larger oscillatory power in ABX mice. (**H)** *Top,* representative 5-s long traces showing unfiltered LFP, as well as filtered theta (4–12 Hz), beta (20–30 Hz), gamma1 (30–50 Hz), gamma2 (50–90 Hz), and gamma3 (90–150 Hz) bands. *Bottom,* representative 5-min long spectrograms. (**I)** *Top,* power spectral density (PSD) averaged across all the analyzed bouts of movement. *Bottom,* bar graphs for the power of theta (\**P* = 0.038, MMANOVA), beta (*P* = 0.173, MMANOVA), gamma1 (\**P* = 0.047, MMANOVA), gamma2 (\*\*\**P* < 0.001, MMANOVA), and gamma3 (\*\*\**P* < 0.001, MMANOVA) bands (*n* = 40 movement bouts from 4 CON, 40 movement bouts from 4 ABX).

We examined grid cells, the predominant cell type in the MEC, whose receptive fields form evenly spaced grids that tile the environment. We generated firing rate maps and spatial autocorrelations of these maps, which showed a clear pattern of spatially correlated spike activity with a center peak and correlation peaks arranged in a hexagonal formation in CON mice (**Fig. 3C**). Grid cells in ABX mice failed to exhibit the characteristic hexagonal patterns (**Fig. 3C**). Quantification of grid-like firing activity revealed significantly lower grid scores, along with significantly lower SI, in ABX mice when compared to the CON group (**Fig. 3D**). Moreover, while ABX grid cells had similar overall MFRs and in-field MFRs compared to CON grid cells, they exhibited significantly higher out-of-field MFRs and a decreased in-field-out-of-field index (**Fig. 3D**). Together, these data indicate that grid cells in ABX mice fire significantly more outside of their receptive field compared to CON mice and have dysfunctional spatial and geometric firing properties. In contrast, other neuron types showed no significant differences between groups, including border cells (**Fig. 3E**), aperiodic spatial cells (**Fig. 3F**), and interneurons (**Fig. 3G**).

We measured the local field potentials (LFPs) in our entorhinal recordings, focusing on the theta (4–12 Hz), beta (20–30 Hz), gamma1 (35–50 Hz), gamma2 (50–90 Hz), and gamma3 (90–150 Hz) frequency bands (**Fig. 3H**). We calculated the power spectral density and quantified the area under the curve for each band during epochs of mouse movement. We found significantly increased power for theta, gamma1, gamma2, and gamma3 bands in ABX mice compared to CON mice, but no significant difference in the beta band (**Fig. 3I**).

### Impaired place cell ensemble function in the hippocampus of ABX mice

To further study the impaired spatial cognition of ABX mice, we investigated place cells in the CA1 field of the hippocampus, which is recognized as a key neural substrate for encoding space. We implanted CON and ABX mice with multielectrode arrays targeted to CA1 (**Fig. 4A**). Firing rate maps were generated from place cells recorded in an open-field arena, which exhibited dispersed firing patterns and larger field sizes in ABX mice compared to CON mice (**Fig. 4B**). We quantified place cell function (by measuring their area and SI) and found significantly larger place field areas and significantly lower SI in ABX place cells (**Fig. 4C**). We also analyzed the firing rates and found that ABX place cells showed significant increases in MFRs and out-of-field MFRs, as well as significantly decreased in-field-out-of-field index but no difference for in-field MFRs when compared to CON place cells (**Fig. 4C**). We calculated the location of the center of mass (COM) of place fields in a subset of our recordings and found that in ABX mice, the location of the COM was significantly closer to the center of the recording chamber (**Fig. 4D**). We compared the COMs across consecutive sessions and found that the shift from one session to the next was significantly greater in ABX mice (**Fig. 4E**). Together, these data indicate dysfunctional CA1 place cells in ABX mice.

**Fig. 4.**
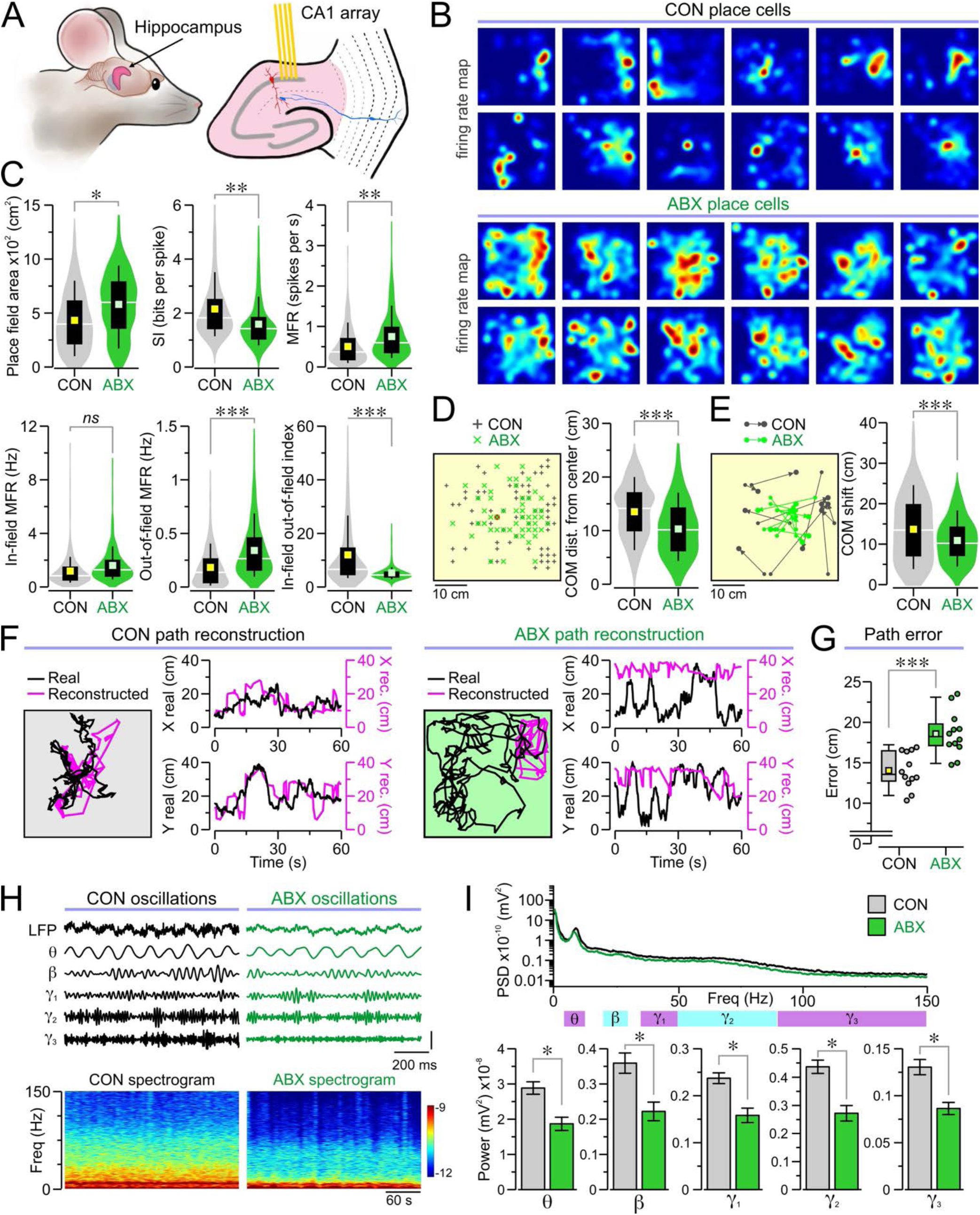
Neurophysiological recordings in the hippocampus. **(A)** Illustration depicting the position of tetrode array placement within the CA1 field of the hippocampus. **(B–E)** Analysis of place cells reveals large disruptions of place cell properties in ABX mice compared to CON mice. (**B)** Representative place cell rate maps in both groups. (**C)** Violin plots of place field area (\**P* = 0.042, MMANOVA), SI (\*\**P* = 0.007, MMANOVA), MFR (\*\**P* = 0.008, MMANOVA), in-field MFR (*P* = 0.087, MMANOVA), out-of-field MFR (\*\*\**P* < 0.001, MMANOVA), and in-field-out-of-field index (\*\*\**P* < 0.001, MMANOVA) for each place cell (*n* = 451 cells from 10 CON, 365 cells from 10 ABX). (**D)** *Left,* map showing the center of mass (COM) for a subset of place cells. *Right,* violin plots of the distances of the COM from the center of the chamber (\**P* = 0.016, MMANOVA; *n* = 372 cells from 7 CON, 195 cells from 6 ABX). (**E)** *Left,* map showing the COM shifts for a subset of place cells. *Right,* violin plots of the COM shifts (\**P* = 0.037, MMANOVA; *n* = 105 cells from 7 CON, 94 cells from 6 ABX). **(F–G)** Place cell ensembles in ABX mice show large reconstruction errors when compared to CON mice. (**F)** Representative real (black) and reconstructed (pink) paths from a 1- min recording interval in CON and ABX mice. Each example shows the real and reconstructed trackplots (*at left*), along with the real and reconstructed X and Y positions (*at right*) during each point in time. **(G)** Box-and-whisker plot (each dot represents a recording interval) of the path reconstruction errors (\*\*\**P* < 0.001, *t* test; *n* = 12 CON sessions, 12 ABX sessions). **(H–I)** Analysis of LFPs during movement bouts reveals much weaker oscillatory power in ABX mice. (**H)** *Top,* representative 5-s long traces showing unfiltered LFP, as well as filtered theta (4–12 Hz), beta (20–30 Hz), gamma1 (30–50 Hz), gamma2 (50–90 Hz), and gamma3 (90–150 Hz) bands. (**I)** *Top,* PSD averaged across all the analyzed bouts of movement. *Bottom,* bar graphs for the power of theta (\**P* = 0.023, MMANOVA), beta (\**P* = 0.024, MMANOVA), gamma1 (\**P* = 0.034, MMANOVA), gamma2 (\**P* = 0.024, MMANOVA), and gamma3 (\**P* = 0.016, MMANOVA) bands (*n* = 100 movement bouts in 10 CON, 100 movement bouts in 10 ABX).

Importantly, the firing activity of place cell ensembles provides a neural code for the animal’s position. We investigated the ensemble function of CA1 place cells with Bayesian path reconstructions, where the activity of a population of place cells is used to reconstruct the position of the mouse over time. This reconstructed position is compared with the real position at each time point (**Fig. 4F**) to obtain the reconstruction error, a useful metric for assessing how well a population of place cells encodes space. Place cell ensembles in ABX mice showed significantly greater reconstruction errors compared to CON mice (**Fig. 4, F and G**), demonstrating that antibiotic treatment disrupts the spatial encoding properties of place cell populations.

We examined the LFPs in the CA1 recordings during periods of movement and calculated the power of each frequency band (**Fig. 4H**). Remarkably, we found that the powers of the theta, beta, gamma1, gamma2, and gamma3 bands were all significantly lower in ABX mice compared to CON mice (**Fig. 4I**).

### ABX mice have dysbiosis and disrupted BBB integrity

We confirmed gut dysbiosis in ABX mice by measuring the ceca and examining the bacterial composition of stool samples taken from CON and ABX mice. Ceca extracted from ABX mice were hyperpigmented, enlarged, and heavier than those from CON mice (**Fig. 5A**). To confirm that antibiotic treatment successfully altered and depleted the gut microbiome, we performed quantitative polymerase reaction (qPCR) on the 16S rRNA gene in stool samples, and 16S RNA sequencing. ABX mice showed shifts in gut microbiome composition at the phylum level, with fewer Bacteroidetes and more cyanobacteria and proteobacteria relative to CON mice (**Fig. 5B**). We identified a 100-fold reduction in 16S rRNA expression in ABX mice compared to CON mice, indicating substantially reduced overall levels of gut bacteria (**Fig. 5C**). Shannon diversity was significantly lower in ABX mice, demonstrating altered composition of the remaining gut microbiome (**Fig. 5C**). Additionally, analysis at the family level revealed that antibiotic treatment substantially reduced the abundance of bacterial families that drive butyrate production (**Fig 5C**).

**Fig. 5.**
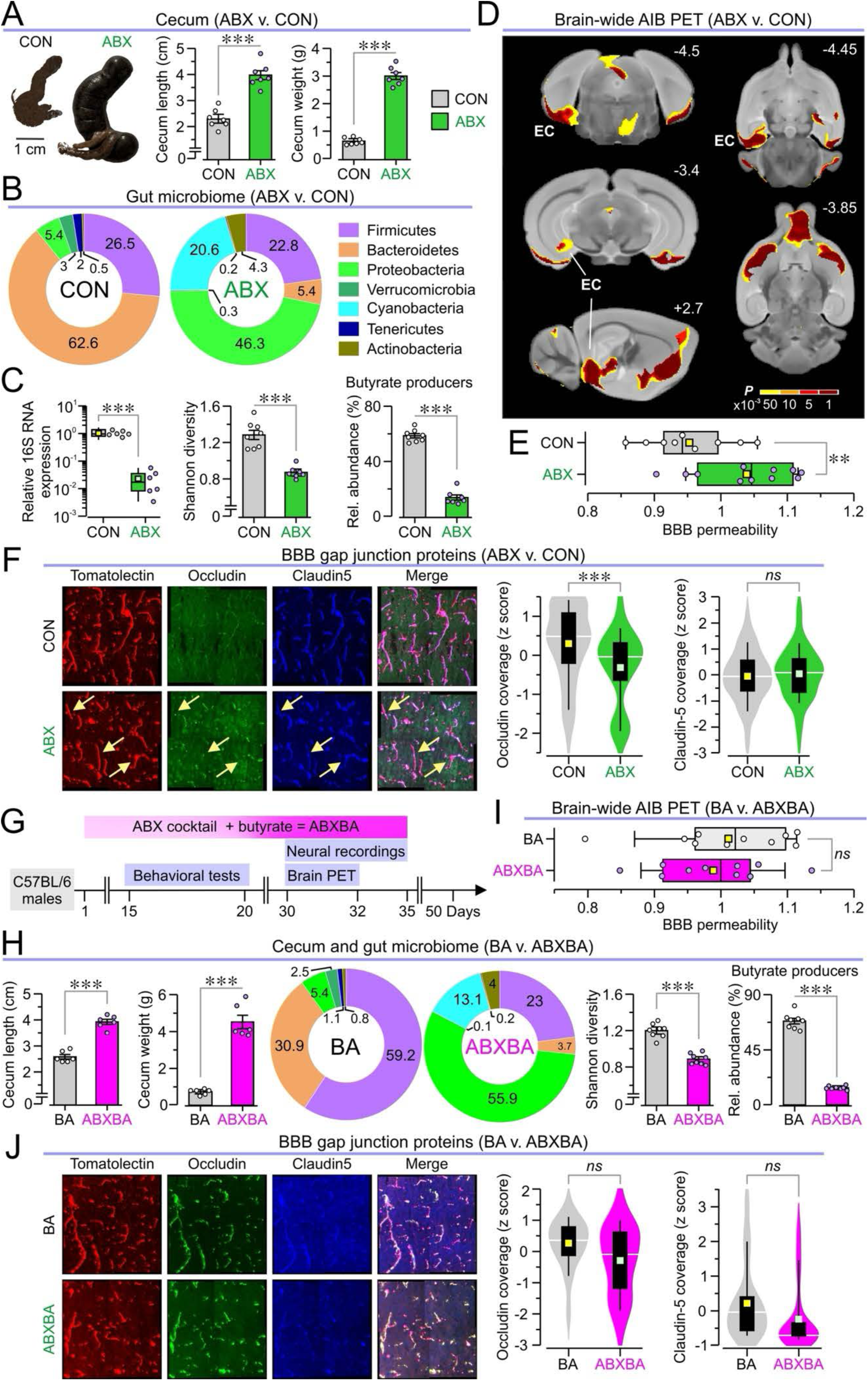
Analysis of the gut microbiome and the effect of butyrate supplementation on ABX mice. (A–C) Antibiotic treatment alters the cecum and depletes the gut microbiome in ABX mice. **(A)** Traced images of ceca (*left*) dissected from CON and ABX mice and bar graphs showing the cecum length (*middle*, \*\*\**P* < 0.001, *t* test) and weight (*right*, \*\*\**P* < 0.001, *t* test) in both groups (*n* = 7 CON, 7 ABX). (**B)** Doughnut plots for the relative abundances of the seven most prominent phyla in CON and ABX mice. (**C)** *Left,* box-and-whisker plot showing the relative expression of 16S RNA (\*\*\**P* < 0.001, *t* test; *n* = 6 CON, 6 ABX); m*iddle,* bar graph of Shannon diversity (\*\*\**P* < 0.001, *t* test; *n* = 8 CON, 7 ABX); *right,* bar graph showing relative abundance of butyrate producers (\*\*\**P* < 0.001, *t* test; *n* = 8 CON, 7 ABX) in stool samples taken from CON and ABX mice. **(D–F)** ABX mice exhibit disruption in the integrity of the blood-brain barrier (BBB) especially around the entorhinal cortex. (**D)** Data of voxel-wide search over the whole brain showing areas in which AIB SUVs are increased in ABX mice using increasing thresholds of significance (scale bar for *P* values <0.001, <0.005, <0.01, <0.05), EC, entorhinal cortex. (**E)** Box-and-whisker plot showing the relative BBB permeability obtained from AIB SUVs in the cluster corresponding to the EC (\**P* = 0.013, *t* test; *n* = 9 CON, 11 ABX). (**F)** *Left,* representative microscopy images staining for tomatolectin, occludin, and claudin-5 in the EC of CON and ABX mice. *Middle,* violin plot of the z-scored coverage of tomatolectin-positive areas with occludin-positive areas (\*\*\**P* < 0.001, MMANOVA). *Right,* violin plot of the z-scored coverage of tomatolectin-positive areas with claudin-5-positive areas (*P* = 0.523, MMANOVA) in CON and ABX mice (*n* = 147 slices in 4 CON, 141 slices in 4 ABX). **(G–J)** Butyrate supplementation does not rescue the gut microbiome composition but restores BBB integrity. (**G)** Experimental timeline for animals in which butyrate is added to the antibiotic cocktail (ABXBA mice); control mice have butyrate added to the drinking water (BA mice). (**H)** *Left,* Bar graphs of cecum lengths (\*\*\**P* < 0.001, *t* test) and weights (\*\*\**P* < 0.001, *t* test) in BA and ABXBA mice (*n* = 6 BA, 6 ABXBA). *Middle,* same as **B** but BA and ABXBA mice. *Right,* bar graphs of Shannon diversity (\*\*\**P* < 0.001, *t* test) and the relative abundance of butyrate producers (\*\*\**P* < 0.001, *t* test) in both groups (*n* = 8 BA, 8 ABXBA). (**I)** Same as **E** but for BA and ABXBA mice (*P* = 0.59, *t* test; *n* = 10 BA, 10 ABXBA). **(J)** Same as **F** but for BA and ABXBA mice, occludin (*P* = 0.101, MMANOVA), claudin-5 (*P* = 0.426, MMANOVA, *n* = 115 slices in 4 BA, 105 slices in 4 ABXBA).

Next, we investigated the BBB in ABX mice. Previous studies have found that germ-free animals, which lack a gut microbiome from birth, possess a leakier BBB, which can be recovered with butyrate treatment (*36*). We addressed this by measuring BBB permeability with PET using the [^11^C]-isoaminobutyric acid (AIB) radiotracer. We identified two regions overlying the entorhinal cortex (bilaterally) with significantly increased AIB uptake in ABX mice, indicating that the BBB was more permeable in the entorhinal cortex (**Fig. 5, D and E**). To confirm that the BBB was compromised, we performed immunohistochemistry, staining for tomatolectin to label blood vessels, and for the tight junction proteins occludin and claudin-5 within the entorhinal cortex. Reductions in the expression of tight junction proteins have been identified in microbiome depleted animals (*36, 16*). We quantified occludin and claudin-5 expression as a percentage of tomatolectin-positive area using automated vessel detection and artifact removal (fig. S3, see Methods for details). After z-scoring areas to account for cross-experiment staining differences, we found that ABX mice had significantly lower occludin coverage, while claudin-5 coverage remained consistent between CON and ABX mice (**Fig. 5F**).

### Butyrate restores BBB integrity but not microbiome composition

Given the virtual disappearance of butyrate producing species in the ABX microbiome (**Fig. 5C**), we investigated whether oral butyrate supplementation could counteract the effects of antibiotic treatment. Mice were treated with butyrate alone (BA mice) or butyrate and the antibiotic cocktail (ABXBA mice), following the same timeline as previous experiments (**Fig. 5G**). We assessed the effects of butyrate supplementation on the gut microbiome. ABXBA mice had significantly elongated and heavier ceca, as well as altered relative abundances of bacterial phyla, decreased Shannon diversity, and a reduction of butyrate producers compared to BA mice (**Fig. 5H**). Together, these results show that butyrate supplementation does not protect the gut microbiome from antibiotic-induced dysbiosis.

We measured BBB permeability with AIB PET, examining the previously identified bilateral cluster in the entorhinal cortex that showed differences between CON and ABX mice. Surprisingly, in this cluster we did not find differences between BA and ABXBA mice (**Fig. 5I**). Moreover, we performed immunohistochemistry, staining for tomatolectin, occludin, and claudin-5, and measured the coverage of occludin and claudin-5 as a percent of the tomatolectin area. Remarkably, there were no significant differences between BA and ABXBA mice for either tight junction protein (**Fig. 5J**). These findings indicate that butyrate supplementation can protect the integrity of the BBB independently of gut dysbiosis.

### Butyrate protects space-encoding neural networks

We sought to determine whether butyrate supplementation may protect spatial cognition during administration of antibiotic cocktail. In the clockmaze task, BA and ABXBA mice spent similar amounts of time, traveled similar distances, and made similar numbers of errors (**Fig. 6A**). Both groups preferentially used the spatial strategy to solve the task, showing similar spatial strategy scores (**Fig. 6B**). During the probe trial, both BA and ABXBA mice spent significantly more time in the target zone surrounding the exit compared to the rest of the maze (fig. S4A). ABXBA mice also demonstrated preserved spatial memory during the OPM task, showing similar investigation times and OPM ratios to those of BA mice (fig. S4B). These results demonstrate that butyrate supplementation preserves spatial cognition despite antibiotic-induced depletion of the gut microbiome. We measured regional brain glucose metabolism in butyrate-supplemented mice with FDG PET and compared SUVs in the previously identified clusters (that showed differences between CON and ABX mice). ABXBA mice had reduced FDG uptake compared to BA mice in the cluster corresponding to the entorhinal cortex, and increased uptake in the cluster corresponding to the anterior hypothalamus (fig. S5A). We then applied our atlas-based analysis and found significantly lower SUVs in ABXBA mice in the lateral entorhinal cortex and MEC when compared to BA mice but not dorsal CA1 (fig. S5B).

**Fig. 6.**
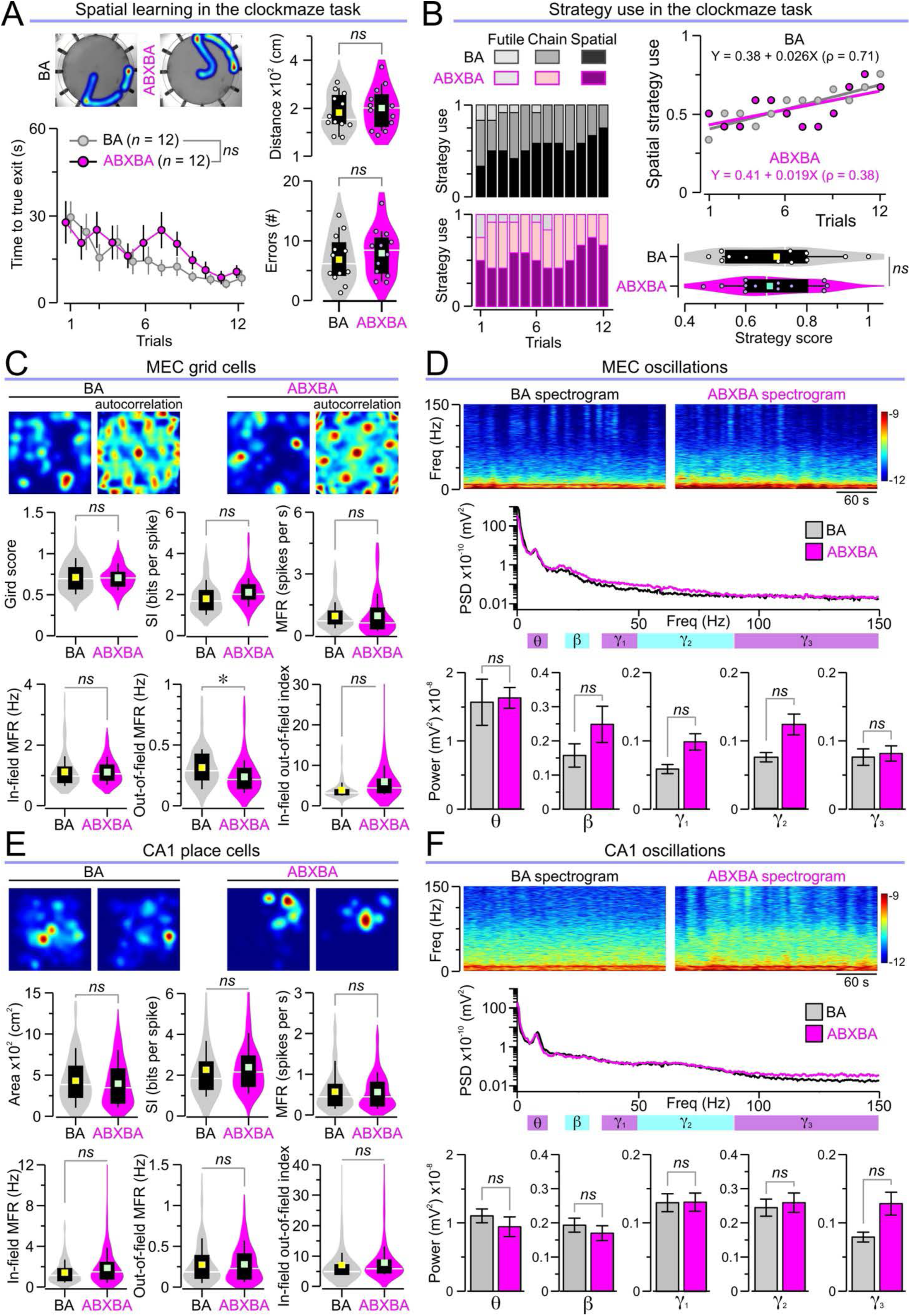
Spatial cognition and neurophysiological recordings in butyrate-treated mice. (A–B) Butyrate supplementation rescues spatial cognition in ABXBA mice subjected to the clockmaze task. (**A)** *Left,* heatmaps of representative paths traveled by mice (*top*) and time series of escape latency (mean ± SEM) showing similar values for BA and ABXBA mice across trials (*P* = 0.59, RMANOVA of log10 transformed times followed by Bonferroni test; *n* = 12 BA, 12 ABXBA). *Right,* violin plots with distances traveled (*top*, *P* = 0.606, *t* test; *n* = 12 BA, 12 ABXBA) and number of errors in all trials (*bottom*, *P* = 0.51, *t* test; *n* = 12 AB, 12 ABXBA). (**B)** ABXBA mice have normal spatial strategy use, as demonstrated by the stacked column graphs with the frequency of the different strategies (*left*), the scatterplot of spatial strategy use (fitted with linear regression curves) for both groups (*top right*) and the violin plots with strategy scores (*bottom right,* \**P* = 0.74, *t* test; *n* = 12 BA, 12 ABXBA) in which higher values reflect higher spatial use. **(C–D)** ABXBA mice have normal grid cells and oscillatory patterns in MEC. (**C)** *Top,* sample grid cell rate maps and corresponding autocorrelations; *bottom,* violin plots of the grid score (*P* = 0.928, MMANOVA), SI (*P* = 0.258, MMANOVA), MFR (*P* = 0.998, MMANOVA), In-field MFR (*P* = 0.89, MMANOVA), out-of-field MFR (\**P* = 0.028, MMANOVA), and in-field-out-of-field index (*P* = 0.088, MMANOVA) for each grid cell recorded from BA and ABXBA mice (*n* = 83 cells in 4 BA, 61 cells in 4 ABXBA). (**D)** *Top,* representative 5-min long spectrogram; *middle,* PSD averaged across all recording bouts analyzed; *bottom,* bar graphs for the power of theta (*P* = 0.837, MMANOVA), beta (*P* = 0.249, MMANOVA), gamma1 (*P* = 0.103, MMANOVA), gamma2 (*P* = 0.124, MMANOVA), and gamma3 (*P* = 0.825, MMANOVA) bands (*n* = 40 movement bouts in 4 BA, 40 movement bouts in 4 ABXBA). **(E–F)** ABXBA mice have normal place cells and oscillatory patterns in CA1. (**E)** *Top,* representative place cell rate maps; *bottom,* violin plots showing place field area (*P* = 0.241, MMANOVA), SI (*P* = 0.427, MMANOVA), MFR (*P* = 0.764, MMANOVA), in-field MFR (\**P* = 0.025, MMANOVA), out-of-field MFR (*P* = 0.92, MMANOVA), and in-field-out-of-field index (*P* = 0.907, MMANOVA) for each place cell in BA and ABXBA mice (*n* = 129 cells in 6 BA, 167 cells in 6 ABXBA). (**F)** *Top,* representative 5-min long spectrogram; *middle,* PSD averaged across all bouts; *bottom,* bar graphs for the power of theta (*P* = 0.591, MMANOVA), beta (*P* = 0.617, MMANOVA), gamma1 (*P* = 0.983, MMANOVA), gamma2 (*P* = 0.807, MMANOVA), and gamma3 (*P* = 0.085, MMANOVA) bands (*n* = 60 movement bouts in 6 BA, 60 movement bouts in 6 ABXBA).

We investigated whether butyrate preserves the neurophysiology of space-encoding neural networks in the entorhinal-hippocampal network. Microelectrode arrays were implanted into the MEC of BA and ABXBA mice, and our unsupervised machine learning pipeline incorporating UMAP and DBSCAN (fig. S6A) identified grid cells, interneurons, aperiodic spatial cells, and border cells (fig. S6, B and C). Remarkably, analysis of grid cell function revealed that BA and ABXBA mice had similar grid scores, SI, MFR, in-field MFR, and in-field-out-of- field indices, though ABXBA mice showed lower out-of-field MFRs compared to BA mice (**Fig. 6C**). Additionally, oscillatory patterns in the MEC LFPs showed no significant differences in the powers of theta, beta, gamma1, gamma2, or gamma3 frequency bands between groups (**Fig. 6D**). We next implanted microelectrode arrays into CA1 in BA and ABXBA mice to assess place cell function. There were no significant differences between groups in place field areas, SI, MFR, in-field MFR, out-of-field MFR, and in-field-out-of-field index (**Fig. 6E**). Analysis of hippocampal LFPs revealed no significant differences between BA and ABXBA mice for theta, beta, gamma1, gamma2, and gamma3 frequency bands (**Fig. 6F**). Together, these neurophysiological results demonstrate that butyrate supplementation preserves the ensemble properties of space-encoding neural networks in both the MEC and hippocampus from the effects of antibiotic treatment.

## DISCUSSION

We demonstrate in this study that oral administration of broad-spectrum antibiotics significantly disrupts the hippocampal–entorhinal network in mice, manifesting in impairments at behavioral, metabolic, vascular, and neurophysiological levels. Specifically, ABX mice exhibit deficits in spatial cognition, as evidenced by their reduced use of spatial strategy in the clockmaze task and impaired place memory in the OPM task. We find decreased glucose metabolism within the MEC, the lateral entorhinal cortex and the dorsal CA1 region of the hippocampus, as revealed by FDG PET. Furthermore, we identify increased BBB permeability localized to the entorhinal cortex in ABX mice, coupled with substantial dysfunction of MEC grid cells –characterized by lower grid scores and aberrant firing patterns– as well as disorganized place cell encoding in hippocampal CA1. Together, these findings underscore a wide-ranging disturbance of the entorhinal-hippocampal spatial coding network following gut microbiome depletion.

Our analysis of gut microbiota confirms the expected state of dysbiosis induced by antibiotic treatment, characterized by decreased microbial diversity, a pronounced reduction in Bacteroidetes abundance and an expansion in proteobacteria, consistent with other reports (*9*, *38*, *41*, *47*). Moreover, dysbiosis leads to a depletion of butyrate-producing bacterial families. Notably, co-administration of butyrate during antibiotic treatment effectively preserved spatial cognitive performance, maintained normal place cell and grid cell function, and sustained BBB integrity, despite persistent microbiome dysbiosis and reduced brain metabolism in some regions. This suggests that butyrate acts as a critical microbiota-derived metabolite that mitigates some of the deleterious neurological consequences of antibiotic-induced microbiome disruption.

Our study introduces an innovative approach to analyzing entorhinal neurophysiology. Current methods for classifying MEC cell types rely on manual inspection or single-metric thresholds, which can be problematic when studying disease models that may alter characteristic firing patterns. To address this limitation, we developed an unsupervised machine learning pipeline that classifies cells based on 11 distinct features. This approach minimizes bias in identifying grid cells and other cell types in both control and experimental conditions, overcoming the limitations of conventional methods that depend solely on grid scores or manual classification. This represents a methodological advance over previous studies of grid cells in disease models, which often rely on potentially biased metrics such as the relative proportion of cells meeting grid score criteria (*72*).

Our results align with previous studies identifying behavioral alterations in microbiome-depleted animals. Previous reports have used both the antibiotic model employed here, or the germ-free model in which animals lack the gut microbiome throughout their lifetime (*48*). These models have found impaired spatial cognition with the Morris water maze task (*14*), addictive behaviors in the conditioned place preference task (*11*, *42*), and impaired extinction learning following a classical cued fear conditioning paradigm (*13*, *49*). Microbiome depleted mice also show altered transcriptomic profiles in the prefrontal cortex and amygdala after fear extinction (*13*, *49*), anatomical changes in the prefrontal cortex, increased neurogenesis in the hippocampus, and changes in dendritic spine density of pyramidal neurons within the amygdala and hippocampus (*51–53*). Neurochemical imbalances in the monoamine levels in the cecum, blood, and brain have also been reported (*47*, *54–58*). Therefore, animals with a depleted gut microbiome have a multitude of anatomic, transcriptomic, metabolic, neurochemical and behavioral alterations that extend beyond the realm of spatial cognition.

Understanding antibiotic-induced adverse effects will help develop therapies that balance pathogen elimination while preserving beneficial commensal bacteria. Various approaches including probiotics, fecal microbiota transfer, narrow-spectrum antibiotics, and CRISPR/Cas9 phagemids have shown promise in mouse models (*59*). Our work suggests that butyrate supplementation could be an effective intervention, though its effects are partial as ABXBA mice remained dysbiotic with reduced entorhinal metabolism. Translating these findings to humans will require optimizing dosing, timing, and delivery methods. Combination therapies, particularly those that restore microbial diversity, may prove highly effective. Given the role of the gut-brain axis in neurological diseases (*18*, *60–63*), including Alzheimer’s disease, Parkinson’s disease, and autism spectrum disorder, understanding how microbiome metabolites influence neural circuits could advance treatments for multiple conditions.

Our study has limitations that should be acknowledged. While we focused on butyrate, other microbial metabolites or immune-modulating factors may also contribute to the observed phenotypes and warrant further investigation. Our antibiotic cocktail, while effective, does not fully mimic the complex dynamics of human antibiotic exposure or natural dysbiosis. Likewise, in humans, spatial cognition is influenced by multifactorial inputs difficult to capture fully in murine models. Lastly, although our PET imaging and immunohistochemical analyses reveal alterations to regional metabolism and BBB permeability, the causal relationships with cognitive impairment require further elucidation using interventional approaches.

In conclusion, this work highlights the vital role of gut microbiome-derived butyrate in preserving the integrity of neural networks that encode spatial cognition and the BBB during antibiotic-induced dysbiosis. By illuminating the pathways through which the microbiome communicates with the brain, our findings advocate for integrative approaches to minimize cognitive side effects of antibiotic treatment and for the potential development of metabolite-based adjunct therapies.

## MATERIALS AND METHODS

### Ethics Statement

Animal experiments followed NIH and ARRIVE guidelines under protocols approved by the Feinstein Institutes for Medical Research Institutional Animal Care and Use Committee (IACUC) (protocol #2023–003). The Feinstein Animal Research Program is registered with the Department of Health and Human Services (DHHS), Office of Laboratory Animal Welfare (OLAW), U.S. Department of Agriculture (USDA #21R0107), Public Health Service (PHS #A3168–01) and New York State Department of Health (NYSDOH #A- 060).

### Animals

Male C57BL/6 mice (3–4 months, Jackson Laboratories, Bar Harbor, ME, USA) were housed at the Center for Comparative Physiology at the Feinstein Institute for Medical Research. Mice were housed in groups of four per cage and were maintained on a reverse schedule of 12 hours of darkness (09:00 to 21:00) and 12 hours of light, with ad libitum access to food and water. Animals were handled over 3 days (15 min per day) before testing. Handling and subsequent experiments were conducted during the dark period of the circadian cycle. Mice with implants for neural recordings were housed individually following surgery.

### Antibiotic treatment

Mice received either normal drinking water (CON), antibiotic cocktail (ABX), butyrate (BA), or antibiotic cocktail supplemented with butyrate (ABXBA) in their drinking water for 2 weeks prior to the start of behavioral testing, and 4 weeks prior to the onset of PET or neural recordings. Animals were maintained on their respective treatment for the duration of experiments. The antibiotic cocktail consisted of ampicillin (0.5 mg per mL, Santa Cruz), metronidazole (0.5 mg per mL, Sigma), neomycin (0.5 mg per mL, Sigma), gentamicin (0.5 mg per mL, Gemini Bioproducts), and vancomycin (0.25 mg per mL, Chem-Impex International), as described previously (*13*). Sodium butyrate (Sigma) was added to the appropriate solutions to reach a concentration of 11 mg per mL (*36*, *64*). Solutions were made fresh and replaced 3 times per week. The body weight of each mouse as well as the food and water intake per cage were measured daily to monitor consumption of the antibiotic solutions. Each cohort was run independently to prevent coprophagia and contamination of the gut microbiome during experiments.

### Cecum measurements

Mice were euthanized with carbon dioxide and cervical dislocation prior to collection of ceca. Animals were placed in a supine position and a midline incision was made to expose the abdominal cavity. The cecum was extracted from the animal by severing the connections to the small and large intestines. Cecum length was measured as the distance between the base, just below the junction between the small and large intestines, and the tip of the cecum. The mass included both the cecum tissue and luminal contents.

### 16S rRNA amplicon sequencing and analysis

Stool samples were collected at 4 weeks of treatment. Animals were placed in a chamber (with grid floor) for 90 min. Autoclaved aluminum foil inserts were placed underneath the grid floor to allow stool collection. Pellets were collected in centrifuge tubes (2 mL), immediately placed on dry ice, and stored at -80°C. Samples were processed in-house for DNA extraction, PCR amplification, and 16S rRNA sequencing. Sequencing was performed for the V3-V4 hypervariable regions using a MiSeq platform (Illumina). Sequencing data were used to calculate Shannon diversity, and relative abundance of taxa at the phylum level. Relative abundance was calculated by dividing the number of reads corresponding to a particular taxon by the total number of reads within a given sample.

### 16S qPCR

DNA was extracted from stool samples collected from mice using MO BIO PowerSoil DNA Isolation Kit (MO BIO Laboratories). Equal amounts of purified DNA were added to qPCR reactions with universal 16S primers (UniF340: 5’-ACTCCTACGGGAGGCAGCAGT-3’; UniR514: 5’-ATTACCGCGGCTGCTGGC-3’) that have previously been validated (*13*). 16S DNA levels in each sample were normalized to the average of the control mouse group.

### Open field task

The apparatus consisted of a square base (40 cm^2^) with 40 cm high walls. The walls and floor of the chamber were painted grey. The floor was covered with a thin layer of bedding that resembled bedding in the home cage. A light bulb (40 W) of orange-red hue illuminated the chamber from above. An infrared-sensitive camera (Marshall Electronics, Torrance, CA, USA) was mounted above the chamber and was connected to the video input of the behavioral tracking software (EthoVision, XT15, Noldus). Mice were subjected to 4 open-field sessions (15 min each) with 2 sessions per day across consecutive days.

### OPM and NOR tasks

The OPM and NOR tasks occurred a day after the last day of open field. OPM utilized the same apparatus as open field. The task consisted of an acclimation trial (T1), a sample trial (T2), and a choice trial (T3). Trials were interspersed with 10 min delay periods, in which each mouse was returned to its home cage. For T1, animals were placed in the empty chamber for 15 min. For T2, mice explored the chamber for 5 min in the presence of two identical objects, which were placed in the center of the NW and NE quadrants, such that each object was 10 cm from the near walls and 30 cm from the distant walls. Four pairs of identical objects of similar size were used. Each object was glued to a metal platform to prevent displacement of the object by the mice. For T3, mice explored the chamber for 5 min with the object in the NE quadrant moved 20 cm to the center of the SE quadrant and was referred to as the ‘moved object’. The object in the NW quadrant was left alone and was referred to as the ‘stable object’. The NOR task began 10 min after the end of T3 and consisted of a second sample phase (T4) and choice phase (T5). During T4, the SE object was returned to its original position and the mouse investigated both objects for 5 min. After another 10-min delay period, the NW object was replaced with a different object and was identified as the ‘novel object’. The NE object was unmanipulated and was referred to as the ‘familiar object’. Mice then underwent T5, in which they spent 5 min investigating the familiar and novel objects. A circular zone around each object was assigned in EthoVision, which recorded the epochs in which the animal’s nose was in proximity (< 1 cm) to the object’s periphery. The number of visits to each object and the total investigation time of the objects were measured. For T2, the investigation ratio was defined as the time exploring the right object (in the NE zone) divided by the sum of the times spent exploring both objects. For T3, the OPM ratio was defined as the time spent exploring the moved object minus the time spent exploring the stationary object divided by the total times exploring both objects. For T5, the NOR ratio was defined as the time spent exploring the novel object minus the time spent exploring the familiar object divided by the total times exploring both objects.

### Clockmaze task

We have previously validated the clockmaze task to assess spatial cognition (*65*), featuring a circular, waterproof base (85 cm diameter) with a clear 30 cm wall, placed on a table in a 4m-by-4m area under video tracking (infrared camera, Marshall). Three illuminated facemasks (illuminance, ∼40 lux) served as distal cues, and water (18±1°C, 2 cm deep) allowed mice to paddle while touching the floor. The perimeter contained 12 equidistant holes (4 cm diameter) fitted with black tubes; 11 were plugged as decoys, and one led to a dry escape pipe (termed ‘true exit’), visually indistinguishable inside the maze. After each trial, mice were dried under a heat lamp. Pre-training over 2 days familiarized mice with escape tasks: Day 1 involved four 60-second trials in a black corridor with an escape tube, and Day 2 involved 4 trials in a clear corridor placed within the clockmaze. Testing on Days 3 and 4 involved 12 trials total, with mice placed in the maze center and 11 plugs acting as decoys, while the single true exit remained constant; mice failing to escape within 60 seconds were gently guided out. Between trials, feces were removed to prevent olfactory cues, and a 15–20 min intertrial interval was used. On Day 5, a probe test with all exits blocked assessed exploratory behavior over 60 seconds. Using EthoVision tracking, we recorded escape latency (time to find the true exit), distance, occupancy paths, and errors—defined as nose or half body inspections into decoy zones (2 cm radius). Escape strategies were classified per trial: spatial (<4 errors within 60 s), chain (>4 errors within 60 s), or futile (no escape within 60 s), with strategy scores derived from these rankings. For analysis of the probe test, the maze was sectioned into twelve equal arcs—one target zone (TZ) containing the true exit and eleven non-target zones (NZ) with false exits—and the time spent in each was measured.

### Other behavioral tasks

The elevated plus maze apparatus was made of grey acrylic elevated 1 m above the ground and consisted of two open arms (35 x 5 cm) and two closed arms (35 x 5 cm). The closed arms had 35 cm tall walls. The light-dark chamber apparatus (Noldus) was constructed of acrylic and consisted of a light section (40 x 20 cm, no overhead cover), and a dark section (20 x 20 cm, infrared-transparent dark acrylic roof). This task was done in bright lighting with an added infrared light (Noldus), and an infrared camera (Marshall Electronics, Torrance, CA, USA) was used to record the mouse. The zero maze apparatus (Maze Engineers, Skokie, IL, USA) consisted of a circular maze with 4 quadrants of alternating open and closed segments. The closed segments had 40 cm tall walls. All three tasks were quantified with ratios calculated as the amount of time spent in the closed (or dark) section minus the amount of time spent in the open (or light) section divided by the total amount of time spent in either section. For the rotarod task, the apparatus (Med Associates, St. Albans, VT, USA) consisted of a five lane rotating drum. Mice underwent five trials with linear acceleration from 4 to 40 rotations per minute over 5 min. Riding time was recorded with Rota Rod 2 (Med Associates) software with a maximum of 5 min per trial.

### Electrode array implantation

Electrode arrays consisted of 4 tetrodes mounted onto custom 3D-printed microdrives, and attached to an Omnetics EIB-16 (Neuralynx, Bozeman. MT). The body of the microdrive was printed using a 3D-printer (Form-3L, Formlabs, Somerville, MA) and the EIB attached to the top using small screws. Tetrodes were created using 17-μm wire (90% platinum, 10% iridium, California Fine Wire, Grover Beach, CA), which was wound so that there were roughly 30–40 rotations per 1 cm length of wire. The four wires were then fused together with a heat gun. Tetrodes were threaded through polyamide tubes, which were glued to a block on the 3D-printed microdrive that could be lowered by turning a calibrated machine screw. After microdrive assembly, tetrodes were electroplated with platinum black solution (Neuralynx) so that impedances were below 350 kꭥ.

For implantation surgeries, mice were anesthetized with 2.5% isoflurane and maintained at 2%, with anesthesia depth regularly assessed by tail or toe pinch reflex. Before starting surgery, the surgical site was prepared by shaving and wiping with betadine solution and isopropyl alcohol. An incision was made along the midline of the scalp, and the overlying connective tissue was removed before covering the skull with Metabond Quick Adhesive Cement (Parkell, Edgewood, NY). A surgical drill (Foredom Electric, Bethel, CT) was used to make a craniotomy above the cerebellum, where the ground screw was inserted into the skull. For surgeries targeting dorsal CA1, a second craniotomy was made on the left side, 2.18 mm posterior to bregma and 1.5 mm lateral from midline. For surgeries targeting the MEC, the second craniotomy was made on the right side, 0.25 mm anterior to the transverse sinus and 3.25–3.5 mm lateral to midline. For MEC implants, tetrodes were inserted at a 3°–5° to the sagittal plane so that the tips were pointed in the posterior direction. The dura was removed using fine forceps, and the tetrodes were slowly lowered to a depth of 0.8 mm below the surface of the brain. Once the tetrodes were in place, the microdrive was fixed to the skull using Ortho-Jet dental acrylic (Lang Dental Inc, Wheeling IL). The ground screw was attached to the EIB-16 using a copper wire, and a 3D- printed cap was glued in place around the microdrive for protection. While the dental acrylic hardened, mice received subcutaneous injections of buprenorphine (Buprenex, 0.05 mg per kg) and saline (0.5 ml). Mice were allowed to recover from anesthesia in a fresh cage on a heating pad and closely monitored for a few days afterward.

For place cell recordings in dorsal CA1, the electrode array of implanted mice was lowered 35–140 μm each day until the tips were located within *stratum pyramidale*. The correct depth was confirmed by the appearance of theta oscillations, the presence of sharp waves and ripple events, and the existence of multiple units with high spike amplitude per tetrode. For grid cell recordings in the MEC, tetrodes were lowered until a final depth of approximately 1 mm was reached.

### Neurophysiological recordings

Neural activity was recorded using a unitary gain headstage preamplifier (HS-18-CNR-LED-MDR50; Neuralynx) connected to the EIB-16 and signals were transferred to a programmable amplifier (Lynx-8, Neuralynx). Signals were collected and stored on a PC running Cheetah (Neuralynx) acquisition software. All tetrodes were referenced to the ground screw in the skull. The position of the mouse was tracked at 30 Hz using a camera mounted on the ceiling above the recording chamber which tracked a red LED on the headstage and saved the position using Cheetah.

### Behavior in implanted mice

For CA1 recordings, implanted mice underwent 1 day of open field testing in which each mouse experienced three 20-min sessions of free exploration within an empty chamber. Delay periods of 15–20 min each were interspersed between the open field sessions. The open field chamber had a square base (40 cm) and 60 cm tall walls built of polyvinyl chloride. The floor of the recording chamber contained no bedding. Animals were transported to the recording room in a covered home cage. The microdrive was connected to the recording system and the mouse was placed in the chamber and allowed to explore freely. Rest periods between behavioral sessions took place in a home cage. For MEC recordings, implanted mice were placed into a square chamber (1 m on the side, 40 cm tall walls) and allowed to roam freely for sessions lasting 15 min each.

### Spike sorting

Neural signals were analyzed offline. Single units were isolated using the automated software Mountainsort, as described previously (*66*). Single units that contained a clear refractory period in their autocorrelograms were categorized as putative pyramidal neurons or interneurons, with the latter excluded from place cell analysis.

### Analysis of neural spike data

Firing rate maps were generated in MATLAB (version R2024a, MathWorks, Inc., Natick, MA) and Neuroexplorer 5 (NEX Technologies, Colorado Springs, CO) by dividing the arena into bins of 2 x 2 cm. For each bin, the total number of spikes was divided by the dwell time. Bins with animal occupancy less than 0.1 s were excluded from analysis. Firing rate maps were smoothed with a 5 x 5 bin Gaussian kernel for CA1 recordings and a 10 x 10 bin Gaussian kernel for MEC recordings. Firing fields were defined as groups of at least 8 contiguous bins that shared an edge and had a firing rate ≥ 20% of the peak firing rate for that unit. The MFR was defined as the number of spikes that occurred during a recording session divided by the total time of that session. The in-field MFR was defined as the average firing rate for all of the bins which were considered to be parts of firing fields. The out-of-field MFR was defined as the average firing rate for all of the bins which were not associated with firing fields. The in-field out-of-field ratio was defined as the in-field MFR divided by the out-of-field MFR.

Spatial information (SI) was calculated by estimating the rate of information I(R|X) between firing rate R and location X with the formula:

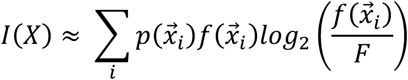

in which p(xi) is the probability for the animal being at location xi, f(xi) is the firing rate measured at location xi, and F is the overall firing rate of the cell.

### Place cell analysis

Place field size was calculated as the area of all of the firing fields for a particular place cell, with fields defined using the 20% peak firing rate described above. If a unit had multiple place fields, then the area of those fields was combined. The COM was calculated as the x and y coordinates corresponding to the weighted average of firing across the columns and rows of the firing rate map. COM distance from center was measured by calculating the distance between the COM of a given place cell and the center coordinate of the chamber for any given session. The COM shift was equal to the distance between the COMs for a given place cell across two consecutive sessions. This metric served as a representation of place field movement across trials.

### MEC single unit analysis

We developed an unsupervised machine learning pipeline to classify units into distinct cell types (fig. S2). The pipeline consisted of feature extraction, normalization, dimensionality reduction, and clustering. For each isolated unit, we generated firing rate maps and spatial autocorrelation maps, from which the metrics to be used as features were calculated. From the firing rate maps, we calculated SI, border score, firing field area, in-field MFR, in-field peak firing rate, out-of-field MFR, and overall MFR. From the spatial autocorrelation maps, we calculated the grid score and performed principal component analysis, selecting the first three principal components as additional features. This feature extraction process was performed separately for two datasets, with the first dataset consisting of cells recorded from the CON and ABX groups, and the second dataset consisting of cells recorded from the BA and ABXBA groups. Extracted features were normalized by calculating z-scores within each dataset to ensure comparable scaling across parameters. The normalized feature vectors were then subjected to UMAP for dimensionality reduction followed by DBSCAN to identify distinct clusters corresponding to different cell types. This unsupervised approach allowed for objective classification of neurons without prior assumptions about cell type boundaries, yielding classifications for grid cells, border cells, interneurons, aperiodic spatial cells, and undefined cells.

### Grid cell analysis

Spatial autocorrelation maps were generated for all classified grid cells using MATLAB scripts adapted from the Moser laboratory. The spatial autocorrelation was calculated as:

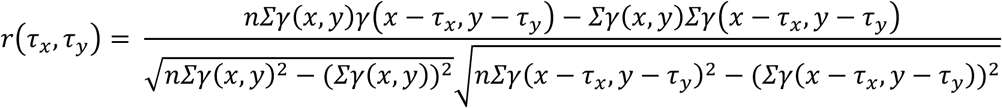

where *γ*(*x*, *y*) is the average firing rate of a cell at location (x,y), and τₓ, τᵧ are spatial lags for which the autocorrelation is calculated. The grid score was determined from the autocorrelogram by calculating the Pearson correlation of the autocorrelation map rotated at angles of 30°, 90°, and 150° versus rotations of 60° and 120°, taking the minimum difference between these two sets of correlations. The grid score metric quantifies the hexagonal rotational symmetry of the autocorrelation maps.

### Border cell analysis

Border scores were calculated according to a previously established method (*67*). Border fields were identified using the firing field definition shown above. The coverage of a given environment wall by a border field was estimated as the fraction of bins along that wall occupied by the field, and cM was defined as the maximum coverage of any single field over any of the four walls of the environment. The mean firing distance dm was computed by averaging the distance to the nearest wall over all bins in the firing rate map belonging to the identified fields, weighted by the firing rate in each pixel. To achieve this, the firing rate was normalized by its sum over all bins belonging to its fields, creating a probability distribution. Then, dm was normalized by half of the shortest side of the environment (representing the largest possible distance to the perimeter) to obtain a fraction between 0 and 1. The border score was calculated as:

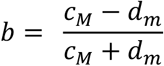

border scores ranged from -1 for cells with central firing fields to +1 for cells with fields that perfectly aligned along at least one entire wall. This measure quantifies the expansion of firing fields along walls rather than away from them, providing an index of border preference, though it saturates when field width approaches half the length of the environment.

### Bayesian Path Reconstruction

A Bayesian two-step approximation with continuity constraint was used to reconstruct the path traveled by the animal (*68*). Recorded position and spike times obtained during open field sessions were imported into MATLAB. Positions were discretized into a 60 x 60 bin grid, with each bin representing .67cm of the chamber. A 10 min encoding period and a 1 min decoding period were used for reconstruction. A sliding window with a width of 1.5 s and a step size of 0.1 s was used. Firing rates for each unit *i* were measured and the map was smoothed with a 2D Gaussian filter with a size of 5 and a standard deviation of 1. The probability for a mouse to be at position *x* given *n* spikes was determined using the following equation:

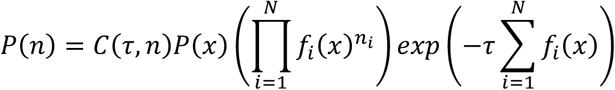

where *f(x)* is the firing rate map, *τ* is the width of the sliding window, and C is a normalizing factor. The estimated position of the mouse was determined as the most probable position generated by the probability distribution:

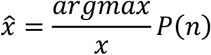

Then, the position probability distribution was calculated for the current time window given the prior using Bayes’ theorem:

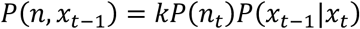

where k is a scaling factor. This allows estimation of the position of the mouse in the current time window:

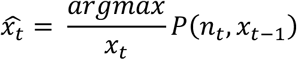

To account for moments of mouse immobility or time windows with a lack of sufficient input information, a continuity constraint based on a Gaussian distribution was applied to the prior using the following formula holding C constant:

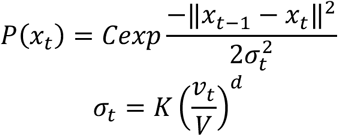

where *σ* relates to how the mouse moves, K and d are constants, K was set to 12, d was set to 1, and V reflects the prior movement speed of the mouse. The reconstruction error was calculated as the distance between the reconstructed position and the real position of the mouse at each time point.

### PET acquisition

Mice underwent PET using the radiotracers [^18^F]-fluorodeoxyglucose (FDG), which assessed glucose metabolism, and [^11^C]-aminoisobutyric acid (AIB) which was used to measure blood-brain permeability throughout the brain (*32*, *69*). Mice were anesthetized with 2.0–2.5% isoflurane in oxygen. After induction, mice were transferred to the scanner where they were maintained under anesthesia with 1.5–2% isoflurane in oxygen. The left lateral tail vein was catheterized with a 30-gauge needle for injection of [^11^C]-AIB (half-life, 20.3 min). Data acquisition began immediately after injection of 1–2 mCi [^11^C]-AIB into the tail vein, using a 60-min dynamic emission scan followed by a 10-min transmission scan. Upon completion of the [^11^C]- AIB scan, 30 min passed to ensure adequate radiotracer decay before IP injections of 1-2 mCi [^18^F]-FDG (half-life, 110 min) with the animals remaining in the same position. Another 40 min passed after the IP injection to allow sufficient uptake of [^18^F]-FDG into the brain, at which point acquisition was initiated. Acquisition was performed for [^18^F]-FDG using a 10-min dynamic emission scan followed by a 10-min transmission scan. Once the scans were completed, animals were allowed to recover in a clean cage and were monitored until the radioactive material decayed.

### PET analysis

Images were preprocessed with PMOD (PMOD Technologies LLC, Zurich, Switzerland) prior to further analysis. The brain portion of each scan was isolated by cropping the acquired images to a box of size 4.9 x 20.7 x 11.9 mm and aligned to an anatomical template generated from magnetic resonance imaging scans of male C57BL/6 mice. Non-brain regions were removed by loading the scans with a brain mask file created from the brain template and using the PMOD manual co-registration functions to eliminate all voxel values outside of the brain space. After PMOD preprocessing, images were analyzed using SPM5 (Wellcome Trust Centre for Neuroimaging, London, UK) with SPM-Mouse (Wolfson Brain Imaging Centre, University of Cambridge, Cambridge, UK; www.spmmouse.org) implemented in MATLAB 2008 (MathWorks, Natick, MA). All images were registered using the realign and reslice function twice, first aligning to a template image and then aligning to the mean of all images. Images were smoothed with an isotropic Gaussian kernel FWHM (full width at half maximum) 0.2 mm at all directions to improve the signal-to-noise ratio.

After pre-processing, whole brain voxel-wise searches were performed using SPM-Mouse for each radiotracer, to identify brain regions in which there were significant differences in uptake between groups. Two sample t-tests were performed across all voxels using a voxel-level threshold of P < 0.001 and a cluster cutoff of 500 voxels to identify significant group differences. The Franklin and Paxinos mouse brain atlas (*70*) was used to identify brain regions and the corresponding clusters were analyzed using FSL (http://www.fmrib.ox.ac.uk/fsl).

The atlas-based approach was performed using MATLAB as previously published (*46*). Masks for all regions-of-interest were generated by drawing templates onto coronal slices for each region based on the Franklin and Paxinos anatomical atlas (*70*). The preprocessed images were loaded into MATLAB and normalized to the mean value for each mouse. Region-of-interest masks were applied to each scan to obtain SUVs for each hemisphere of each coronal slice. Heatmap images were created in MATLAB using the JET colormap, with SUVs for each region overlaid onto corresponding coronal slices downloaded from the Allen Brain Map Reference Atlas (http://atlas.brain-map.org/) for visualization purposes.

### Immunohistochemistry

Mice were anesthetized intraperitoneally using euthasol (100 uL) and transcardially perfused with 0.9% saline containing 0.5% sodium nitrite and heparin 10 units per ml. Brains were removed, embedded in O.C.T., flash frozen on methylbutane at -40°C, and cryosectioned at a thickness of 14um. Sections were thawed for 20 minutes at room temperature and were post-fixed by immersion in ethanol for 30 minutes, followed by acetone for 1 minute. Sections were washed (2x) in 1x PBS before being blocked with 3% BSA (Bovine Serum Albumin) and 0.3% Triton-X in 1x PBS for 1 hour at 4°C. Sections were then incubated with primary antibody overnight at 4°C and were washed (3x) with 0.2% Tween in PBS before incubation with fluorophore-conjugated secondary antibodies for 1 hour. After incubation, sections were washed (3x) with 0.2% Tween in PBS and stained with DAPI (0.5 ug/ml in 1x PBS). Finally, sections were mounted with DAKO- mounting medium (#S3023), and cover-slipped (Fisher). Biotinylated tomato lectin primary antibody (Vector Laboratories, B-1175, 1:200 dilution) was used to stain vascular endothelium (66). For staining tight junction proteins, primary antibodies targeting claudin-5 (Invitrogen, 34-1600, rabbit, 1:100 dilution) and occludin (Life Technologies, 331511, mouse, Occludin-FITC, 5 ug/ml) were used. Secondary antibodies were all prepared using a 1:200 dilution and included Streptavidin Alexa Fluor 594 (Life Technologies, S11227), goat-anti-rabbit Alexa Fluor 647 (Invitrogen, A21245), and goat-anti-mouse AlexaFluor 488 (Invitrogen, A11001). All dilutions for primary and secondary antibodies were performed using 3% BSA in 1x PBS.

### Fluorescent Microscopy Analysis

All coverslipped mounted fluorescent tissue sections were scanned using an Apotome microscope (Zeiss). Images were acquired using Zen Pro software (Zeiss) with 63x oil immersion lens. The rhinal fissure was used as a landmark to identify the entorhinal cortex. Z-stacked tiled 2x2 images (6 stacks, 0.50 um each) were acquired for the entire entorhinal cortex in each section. Each image was processed for analysis by stitching tiles with 10% overlap and compressing z-stacks into a single image using maximum projection.

### Blood vessel mask generation for tight junction protein quantification

To determine the percent coverage of occludin and claudin-5 throughout blood vessels, we created masks of the tomatolectin stain using a six-step automated image processing pipeline in MATLAB (fig. S3): [1] raw grayscale images of tomatolectin immunofluorescence were acquired; [2] artifacts likely corresponding to aggregated antibody were identified by detecting circular objects significantly brighter than their immediate surroundings, as these were clearly distinct from the elongated morphology of blood vessels; [3] these artifacts were removed through interpolation of surrounding pixel intensities, and the resulting image was smoothed using a two-dimensional Gaussian kernel to reduce noise; [4] edge detection algorithms were applied to convert the grayscale image to a binary black-and-white image, with previously identified artifact locations excluded from edge detection; [5] edges were closed and dilated creating continuous vessel boundaries that were subsequently filled through interpolation, and [6] small objects were removed to eliminate residual image noise, generating the final binary mask used for quantifying tight junction protein coverage. This mask was applied to the occludin and claudin-5 images, and the percent coverage of occludin and claudin-5 was then calculated as the ratio of positive immunofluorescence area to total masked area.

### Statistical Analysis

Statistical testing was carried out using Origin Pro (OriginLab, Northampton, MA, USA) and the Statistics and Machine Learning Toolbox in MATLAB. Nested datasets were analyzed with mixed model ANOVA (MMANOVA). The Shapiro-Wilk test was used to determine whether samples were normally distributed. For comparisons between groups, Student’s t-test was used for normally distributed samples and the Mann-Whitney U-test was used for samples that were non-normally distributed. One-Way ANOVA or repeated measures ANOVA (RMANOVA) were used with Bonferroni test or Tukey post-hoc test. Violin plots were fitted with the kernel density estimation. Box-and-whisker plots show the median (horizontal line), mean (square), 25–75% quartiles (box) and 10–90% range (whiskers). Bar graphs show the mean ± standard error of the mean. For bar graphs, violin plots, and box-and-whisker plots with dots, each dot represents one mouse.

## Acknowledgments

We thank Elvira Strohl for the illustrations in Fig. 3 and Fig. 4; Anette Lee for help with the 16S RNA analysis; Pedro Gómez and Joseph Gallagher for help with behavioral assessments; as well as Philippe Marambaud, An Vo, and Stavros Zanos for help in JMG’s MD/PhD thesis committee.

## Funding

National Institutes of Health grant 5P01AI102852 (to PH); National Institutes of Health grant 5P01AI073693 (to PH); Department of Defense (DOD) impact award W81XWH1910759 (to PH); Department of Defense (DOD) impact award HT9425-24-1-0976 (to PH)

## Author contributions

Conceptualization: JMG, JJS, PTH; Methodology: JMG, JJS, JC, CBM, PTH; Investigation: JMG, JJS, JC, CBM, PTH; Visualization: JMG, JJS, PTH; Supervision: PTH; Writing—original draft: JMG, JJS, PTH; Writing—review & editing: JMG, JJS, PTH.

## Competing interests

Authors declare that they have no competing interests.

## Data and materials availability

All data needed to evaluate the conclusions in the manuscript are available in the main text and/or the Supplementary Materials.

## Supplementary Materials

**Supplemental Fig. 1.**
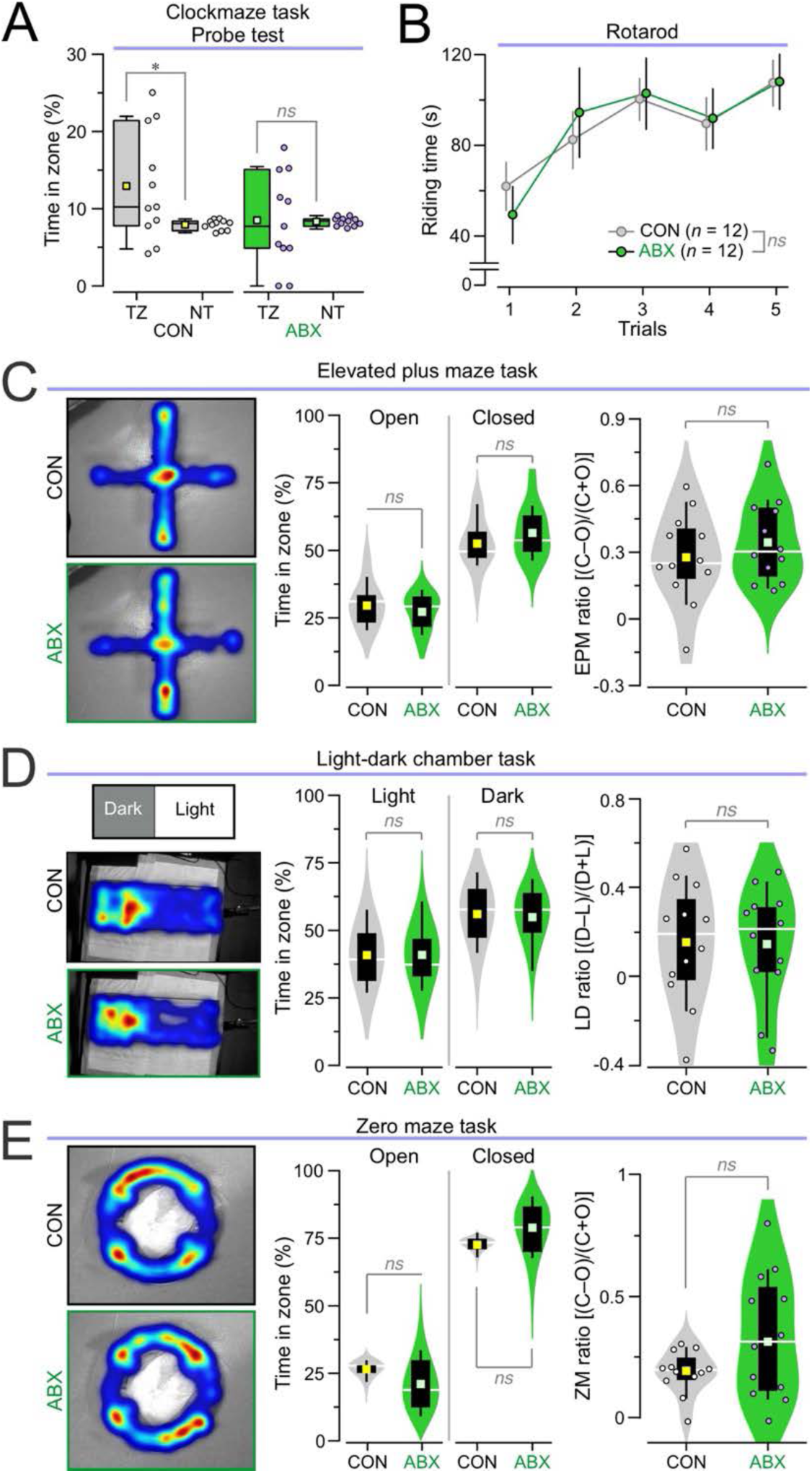
Behavioral tasks in CON and ABX mice. **(A)** Probe test in the clockmaze task: Box-and-whisker plots for the percent of time in the target zone (TZ) or the non-target zone (NZ). CON mice spend significantly more time in the TZ than the NZ (\**P* < 0.031, *t* test, *n* = 11) whereas ABX mice do not show significant difference between zones (*P* = 0.925, *t* test, *n* = 11). (**B)** Rotarod task: dot and line plot showing no differences in riding time across 5 trials in CON and ABX mice (*P* = 0.949, RMANOVA followed by Bonferroni test; *n* = 12 CON, 12 ABX). (**C)** Elevated plus maze (EPM) task: *Left,* representative occupancy heatmaps; *middle,* violin plots with the percent time spent in either the open (*P* = 0.436, *t* test) and closed arms (*P* = 0.305, *t* test); *right,* violin plot with the EPM ratio (*P* = 0.394, *t* test), defined as the time in the closed arms minus the time in the open arms divided by the summed time in both arms, for both groups (*n* = 12 CON, 12 ABX). (**D)** Light-dark (LD) chamber task: *Left,* occupancy heatmaps; *middle,* violin plots with the percent time spent in either the light (*P* = 0.98, *t* test) or dark (*P* = 0.836, *t* test) zones; *right,* violin plot showing the LD ratio (*P* = 0.931, *t* test), defined as the time in the dark zone minus the time in the light zone divided by the time in both zones, for both groups (*n* = 12 CON, 12 ABX). (**E)** Zero maze (ZM) task: *Left,* occupancy heatmaps; *middle,* violin plots with the percent time spent in either the closed (*P* = 0.07, *t* test) or open (*P* = 0.083, *t* test) zones; *right,* violin plot showing the ZM ratio (*P* = 0.071, *t* test), defined as the time in the closed zone minus the time in the open zone divided by the time in both zones, for both groups (*n* = 12 CON, 12 ABX).

**Supplemental Fig. 2.**
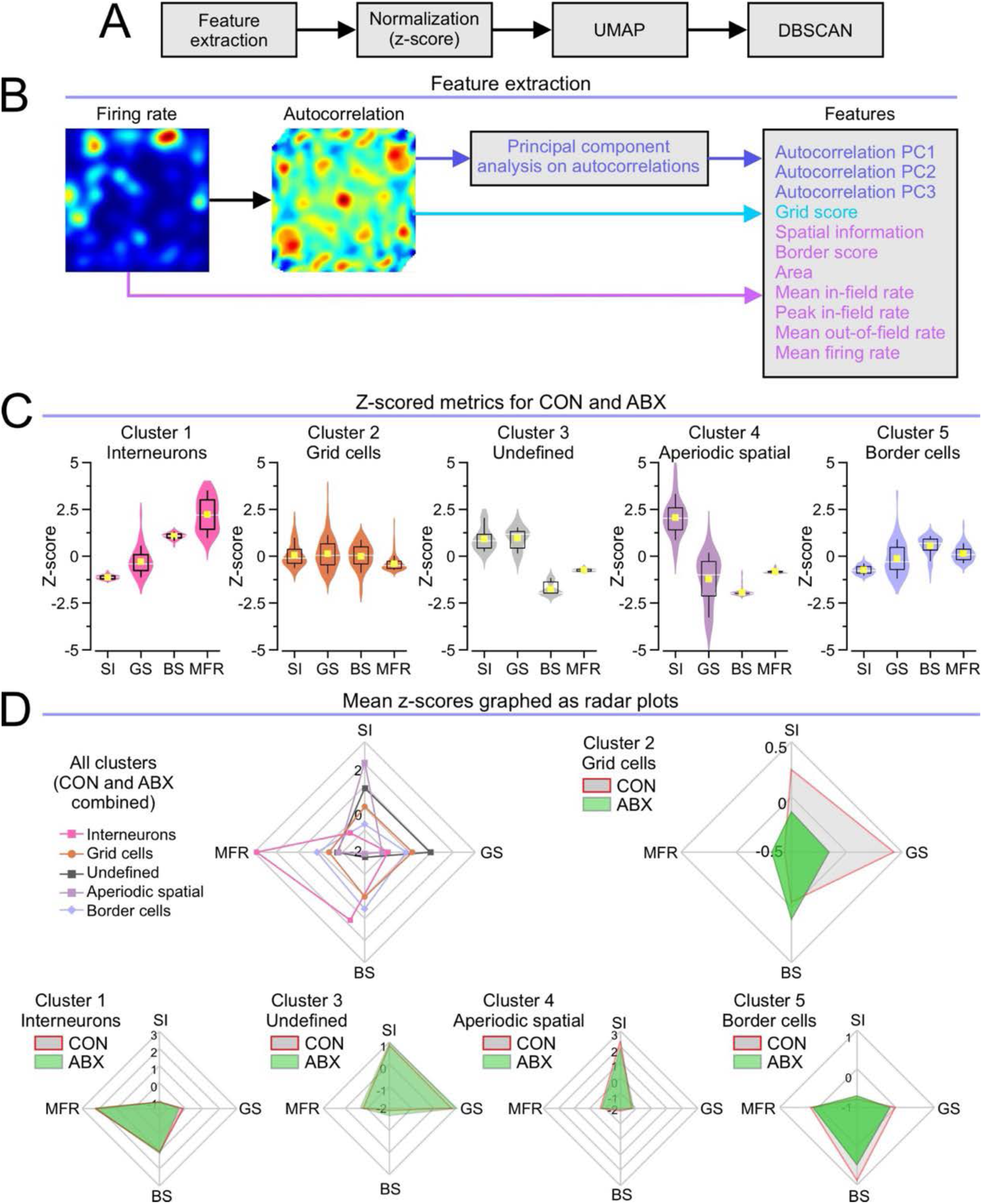
Medial entorhinal cortex neurophysiology analysis pipeline. **(A)** Diagram outlining major steps in unsupervised learning pipeline. For each isolated neuron, features are extracted and normalized by taking the z-score for each parameter. These normalized features are inputted into a UMAP for dimensionality reduction. The dimensionality reduced data is then clustered using DBSCAN to classify each cell type. (**B)** Diagram explaining feature extraction. Firing rate maps and autocorrelations are generated for all cells. From the rate map, we calculate the following features: SI, border score, area, mean in-field rate, peak in-field rate, mean out-of-field rate, and MFR. From the autocorrelation, we calculate the following features: grid score and the first three components of the principal component analysis. This pipeline is applied separately for two datasets (dataset 1: CON and ABX; dataset 2: BA and ABXBA). All of the metrics above are normalized, and the z-scores are used for the rest of the analysis. (**C)** Violin plots showing z-scored metrics for cluster classification. Distinct clusters emerge in each dataset, characterized by their defining features. They are identified as: interneurons (cluster 1, high MFR, low SI), grid cells (cluster 2, high grid score), undefined (cluster 3, mid-range SI and grid score, low MFR and border score), aperiodic spatial (cluster 4, high SI, low grid score), and border cells (cluster 5, high border score). (**D)** Radar plots showing the mean SI, grid score, border score, and MFR for each cluster. In the top left plot, the means are shown for each cluster with CON and ABX mice combined. The remaining plots show each cluster segregated into CON and ABX mice. Remarkably, cluster 2 for grid cells demonstrates that CON and ABX groups can be easily discriminated.

**Supplemental Figure 3.**
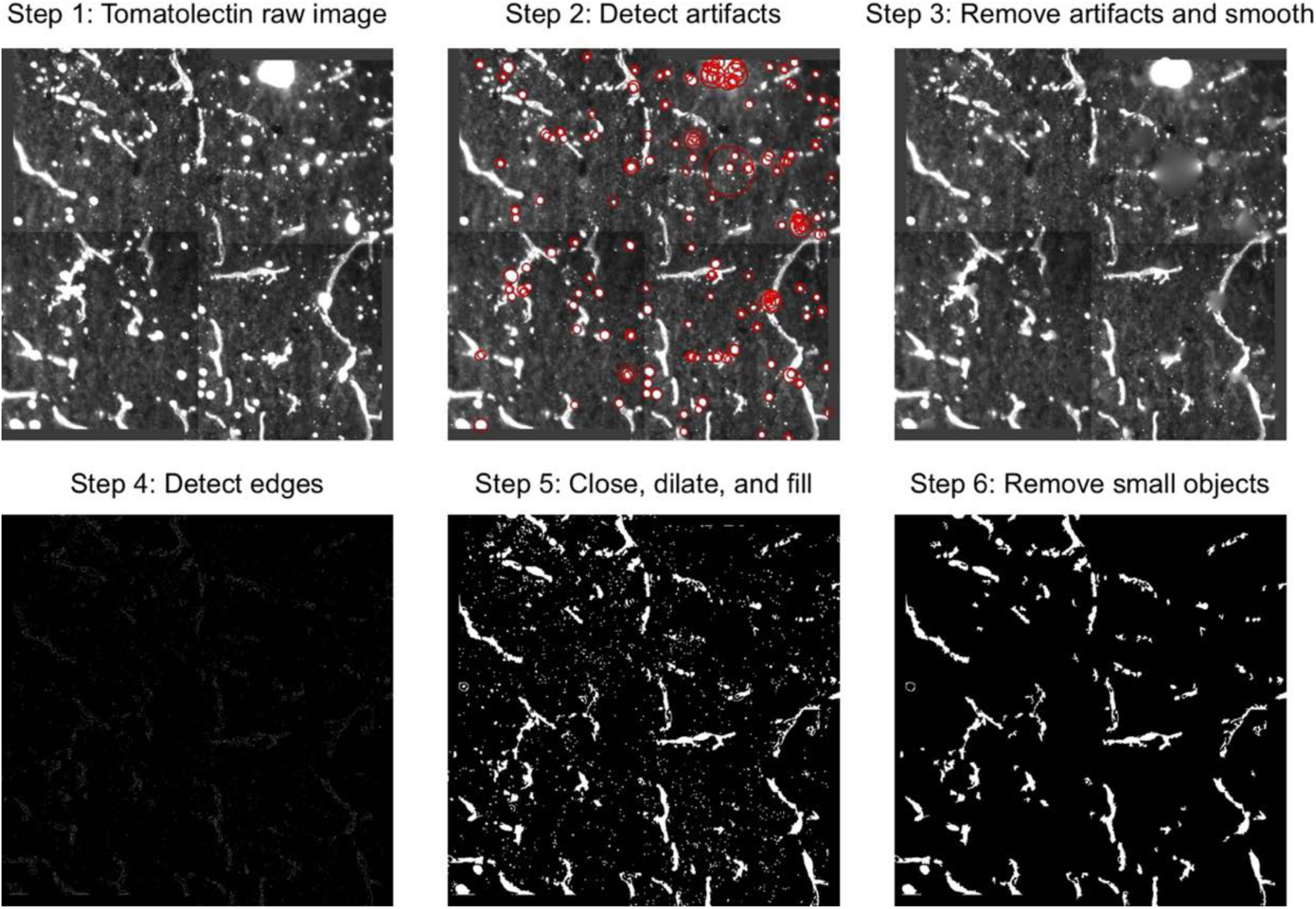
Image processing for blood vessel analysis. To determine the percent coverage of occludin and claudin-5 throughout the blood vessels, we create a mask of the tomatolectin stain with a six-step process. In **step 1**, the raw greyscale image of the tomatolectin stain is acquired. In **step 2**, we detect artifacts which likely correspond to aggregated antibody and are clearly distinct from blood vessels by searching for circular objects which are brighter than their surroundings. In **step 3**, these artifacts are removed through interpolation of the surrounding areas, and the image is smoothed with a 2-dimensional Gaussian kernel. In **step 4**, the edges are detected, excluding those corresponding to the previously identified artifacts, converting the greyscale image to a binary black-and-white image. In **step 5**, the blood vessel outlines are defined using a series of morphological dilation and erosion operations (which respectively expand and shrink the detected edges to create continuous vessel boundaries). These outlines are then filled through interpolation. In **step 6**, objects smaller than 1000 pixels are removed to account for image noise, generating the final mask that is used for subsequent analysis.

**Supplemental Figure 4.**
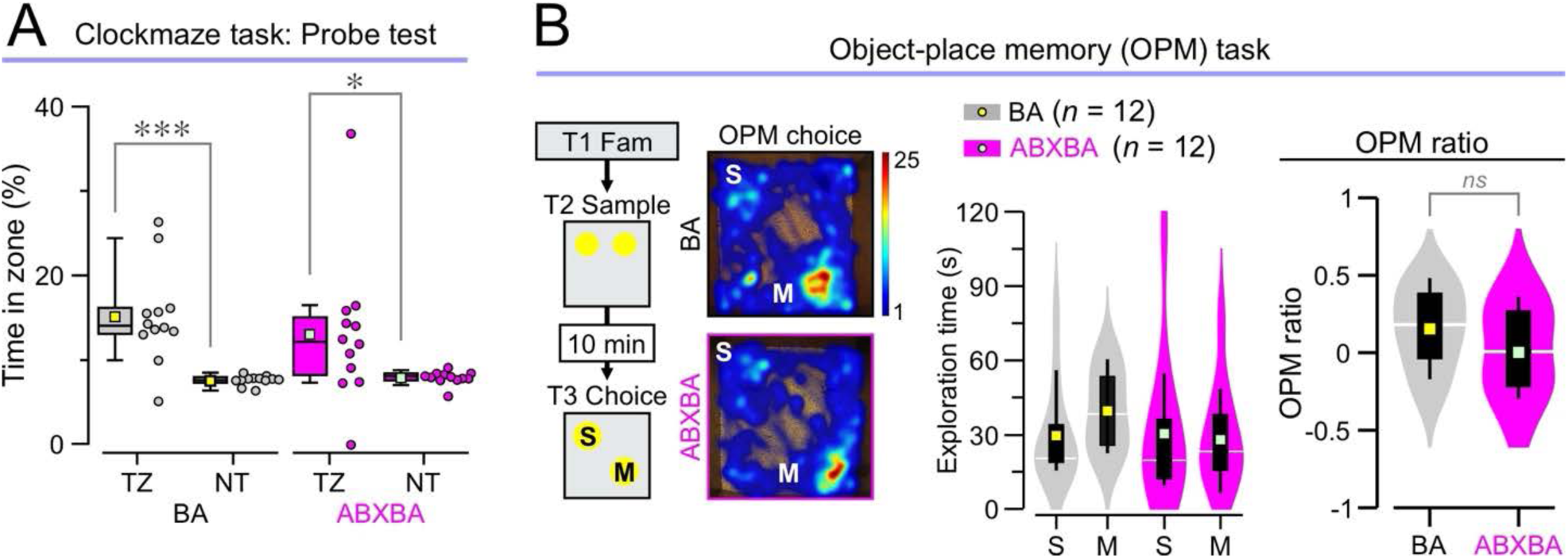
Behavioral tasks in BA and ABXBA mice. **(A)** Probe test in the clockmaze task: Box-and-whisker plots for the percent of time spent in the target zone (TZ) or the non-target zone (NZ). Both groups spend significantly more time in the TZ than the NZ (BA: \*\*\**P* = 0.00059, Mann-Whitney U test, *n* = 12; ABXBA: \**P* < 0.035, Mann-Whitney U test, *n* = 12). (**B)** *Left,* schematic of the OPM task and representative occupancy heatmaps. *Middle,* violin plots showing the investigation times for the stable and moved objects (BA v. ABXBA, *P* = 0.404; stable v. moved, *P* = 0.561, two-way ANOVA followed by Tukey test; *n* = 12 BA, 12 ABXBA). *Right,* Violin plots of the OPM ratios (*P* = 0.234, *t* test; *n* = 12 BA, 12 ABXBA).

**Supplemental Figure 5.**
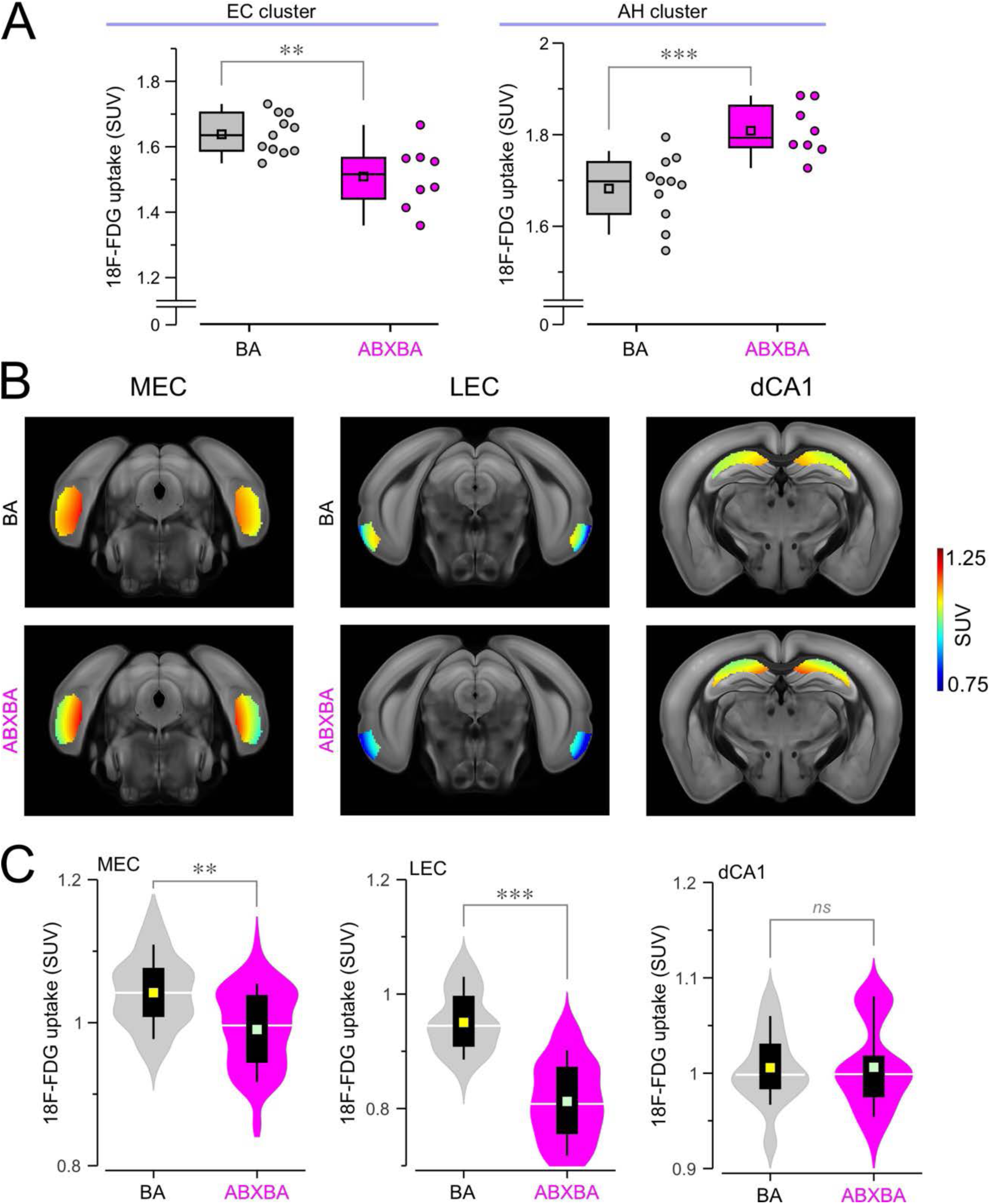
Positron emission tomography in butyrate-treated mice. **(A)** Box-and-whisker plots showing the FDG SUVs for the clusters corresponding to the entorhinal cortex (EC) (*left*) and the anterior hypothalamus (AH) (*right*). Cluster coordinates correspond to those described for CON and ABX mice (Fig. 2, A and B). **(B)** Brain atlas-based analysis with masked slices for the medial entorhinal cortex (MEC), lateral entorhinal cortex (LEC) and dorsal CA1 (dCA1). The heatmaps show the SUVs for one slice averaged across all mice and overlaid onto an MRI template. **(C)** Violin plots showing that ABXBA mice display significantly decreased FDG uptake in the MEC (\*\**P* = 0.001, MMANOVA; *n* = 330 slices from 11 BA, 240 slices from 8 ABXBA), LEC (\*\*\**P* < 0.001, MMANOVA; *n* = 220 slices from 11 BA, 160 slices from 8 ABXBA), and dCA1 (*P* = 0.944, MMANOVA; *n* = 330 slices from 11 BA, 240 slices from 8 ABXBA).

**Supplemental Figure 6.**
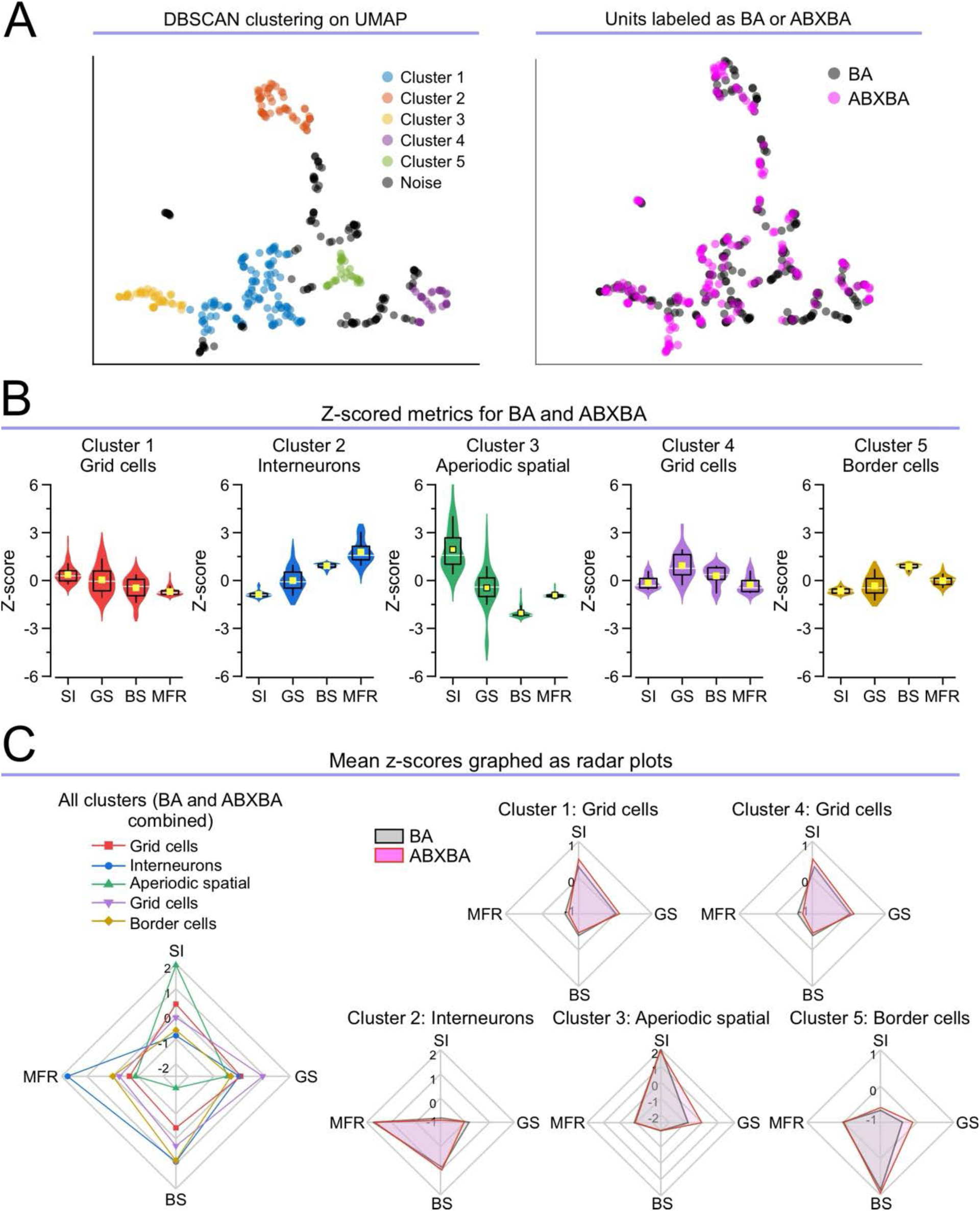
Medial entorhinal cortex neurophysiology analysis pipeline in butyrate-treated mice. **(A)** *Left,* UMAP with DBSCAN clustering. *Right,* UMAP showing group (BA or ABXBA) for each unit. **(B)** Violin plots showing z-scored metrics for cluster classification. The clusters are identified as: grid cells (cluster 1, high grid score, most populous cluster), interneurons (cluster 2, high MFR, low SI), aperiodic spatial (cluster 4, high SI), grid cells (cluster 4, high grid score), and border cells (cluster 5, high border score). **(C)** Radar plots showing the mean SI, grid score, border score, and MFR for each cluster. In the leftmost plot, the means are shown for each cluster with BA and ABXBA mice combined. The remaining plots each show one cluster comparing BA and ABXBA mice.

## Notes

### Competing Interest Statement

The authors have declared no competing interest.

